# MYC enhancer invasion promotes prognostic cancer type-specific gene programs through an epigenetic switch

**DOI:** 10.1101/2023.01.06.522970

**Authors:** Simon T. Jakobsen, Rikke AM. Jensen, Tina Ravnsborg, Christian D. Vaagenso, Hjorleifur Einarsson, Maria S. Madsen, Robin Andersson, Ole N. Jensen, Rasmus Siersbæk

## Abstract

The transcription factor MYC is overexpressed in most cancers, where it drives multiple hallmarks of cancer progression. MYC is known to promote oncogenic transcription by binding to active promoters. In addition, MYC has also been shown to invade distal enhancers when expressed at oncogenic levels, but this enhancer binding has been proposed to have low gene-regulatory potential. Here, we demonstrate that MYC enhancer binding directly promotes cancer type-specific gene programs predictive of poor patient prognosis. MYC induces transcription of enhancer RNA through recruitment of RNAPII, rather than regulating RNAPII pause-release as is the case at promoters. This is mediated by MYC-induced H3K9 demethylation by KDM3A and acetylation by GCN5, leading to enhancer-specific BRD4 recruitment through its bromodomains, which facilitates RNAPII recruitment. Thus, we propose that MYC drives prognostic cancer type-specific gene programs by promoting RNAPII recruitment to enhancers through induction of an epigenetic switch.

## Introduction

MYC is overexpressed in most cancers (Vita & Henriksson, 2006), making it one of the most prominent oncoproteins described to date. A multitude of alterations can result in MYC over-expression, including amplification of the locus (Beroukhim et al., 2010), disruption of upstream signaling pathways (Kress, Sabò, & Amati, 2015), gained enhancer and super-enhancer activity near *MYC* (Affer et al., 2014), and increased protein stability (Junttila & Westermarck, 2008). Overexpression of MYC is known to drive cell transformation by regulating multiple hallmarks of tumorigenesis, e.g., proliferation, metabolism, and invasion (Meškytė, Keskas, & Ciribilli, 2020).

Efforts to understand the mechanisms underlying MYC function in transcriptional regulation have focused on gene promoters, where MYC binds ubiquitously as a heterodimer with MAX (Lee et al., 2012). At gene promoters, MYC has been shown to activate transcription by promoting the transition of paused RNA polymerase II (RNAPII) characterized by serine 5 phosphorylation to its elongating form characterized by Serine 2 phosphorylation (Eberhardy & Farnham, 2002; Lin et al., 2012; Rahl et al., 2010).

Interestingly, MYC seems to affect tumorigenesis in a cancer type-specific manner (Green et al., 2016; Ji et al., 2011; Qiu et al., 2022; Zeid et al., 2018), which is hard to reconcile with the ubiquitous nature of MYC binding to promoters. Interestingly, it has been shown that MYC also starts to invade active enhancer regions upon overexpression (Lin et al., 2012; Sabò et al., 2014; See, Chen, & Fullwood, 2022; Walz et al., 2014; Zeid et al., 2018). It has been suggested that MYC first saturates high-affinity E-box binding sites in promoter regions before invading promoter-distal regions with an alternative low-affinity E-box upon overexpression (Lin et al., 2012; Zeid et al., 2018). This enhancer invasion by MYC has been proposed to have low gene regulatory potential (Walz et al., 2014), and the functional significance of MYC enhancer binding is therefore still elusive.

In this study, we demonstrate that MYC regulates the activity of cancer type-specific enhancers to promote cancer type-specific oncogenic gene programs predictive of patient prognosis. We show that MYC activates enhancer transcription by recruiting RNAPII rather than increasing enhancer RNAPII pause-release. This is mediated by a MYC-activated enhancer-specific epigenetic switch involving the combined action of the general control non-derepressible 5 (GCN5) acetyltransferase and the lysine demethylase 3A (KDM3A). This switch promotes BRD4 enhancer recruitment through its bromodomains, which drives RNAPII recruitment and enhancer transcription.

## Results

### Oncogenic MYC invades cancer type-specific enhancers

To investigate the role of MYC at enhancers, we focused on four breast cancer cell lines representing the two diverse triple-negative (TNBC) and estrogen receptor α-positive (ER+) breast cancer subtypes. Breast cancer cell lines showed intermediate levels of MYC overexpression compared to other solid tumor cell lines (Suppl. Fig. 1A-B) and therefore represent relevant models for studying the role of MYC enhancer invasion in cancer cell biology. Consistent with previous findings (Lin et al., 2012; Sabò et al., 2014; See et al., 2022; Walz et al., 2014; Zeid et al., 2018), chromatin immunoprecipitation coupled to high throughput sequencing (ChIP-seq) revealed that >70% of MYC binding sites were within promoter distal regions across all four breast cancer cell lines (Fig. 1A). Furthermore, MYC peaks were preferentially located at strong E-box motifs at the promoters compared to weaker E-box motifs at the enhancers (Suppl. Fig. 1C) as previously described (Lin et al., 2012; Zeid et al., 2018). Consistently, MYC binding was lower at enhancers compared to promoters in all four breast cancer cell lines (Suppl. Fig. 1D). Interestingly, 82% of these distal binding sites (excluding promoters, exons, and UTRs to avoid unannotated TSSs) overlapped with ENCODE SCREEN enhancers (high DNase and H3K27ac signal) from their consortium of cell lines (Fig. 1B) (Moore et al., 2020). This indicates that gene distal MYC binding sites represent mostly active enhancers consistent with the notion that MYC preferably binds at transcriptional active genomic regions (Guccione et al., 2006; Nie et al., 2012). We will therefore henceforth refer to promoter distal MYC binding sites as enhancers.

**Figure 1.**
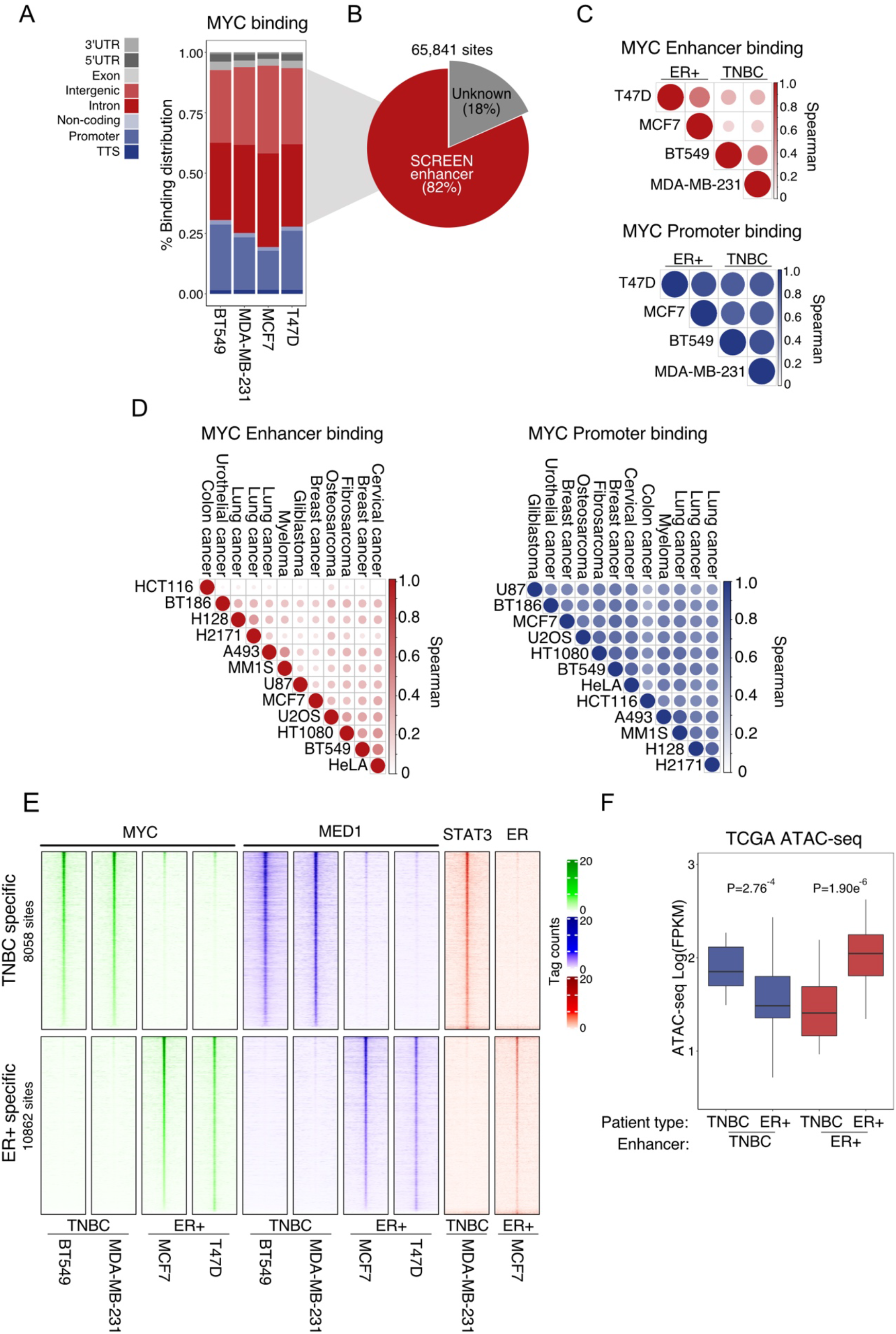
MYC binds cancer type-specific enhancers. **(A)** Genomic annotation of MYC ChIP-seq binding sites in BT549, MDA-MB-231, MCF7, and T47D cells. Peaks were annotated using homer software (Heinz et al., 2010) with the hg38 RefSeq reference genome. Promoters were defined as -1kb to +100bp around the TSS. **(B)** Overlap of promoter distal MYC binding regions (i.e., binding regions outside promoters and exons) with Encode SCREEN human enhancer database (Moore et al., 2020). SCREEN candidate enhancers were defined as regions with high DNase-seq signal and H3K27ac ChIP-seq signal across cell lines in Encode. **(C)** Spearman correlation of MYC binding across the four cell lines in MYC-bound enhancers (n=89,631) or promoters (n=10,049) called in at least one of the four breast cancer cell lines. **(D)** Similar analysis as in panel C, but for 12 cancer cell lines using publicly available MYC ChIP-seq data (GSE86504 (G. Wu et al., 2017), GSE86511 (Galardi et al., 2016), GSE78064 (Xiong et al., 2017), GSE44672 (Walz et al., 2014), GSE36354 (Lin et al., 2012)). **(E)** MYC, MED1, ER (GSE23893 (Joseph et al., 2010)), and STAT3 (GSE152203 (Conway et al., 2020)) binding as determined by ChIP-seq at breast cancer subtype-specific MYC enhancer sites (DESeq2, Padj<0.05 in both cell lines representing each subtype). Data is represented in a heatmap +/-5kb from the center of the peak. **(F)** Accessibility of MYC breast cancer subtype-specific enhancers from panel E in TCGA ATAC-seq data was plotted for TNBC (n=32) and ER+ breast cancer (n=117) patients. The boxplot shows the average Log scaled FPKM counts of the given regions for patients in each group. The data was extracted from (Corces et al., 2018). P-values were calculated using Welch’s two-sided t-test.

A comparison of MYC binding between cell lines revealed that MYC commonly binds the same promoters but cell line-specific enhancers (Fig. 1C). Similar results were obtained from 12 cancer cell lines representing different cancer types (Fig. 1D), indicating that this is a general feature of MYC binding to chromatin in cancer. This is consistent with previous observations for MYCN in neuroblastoma (Zeid et al., 2018). Importantly, breast cancer subtype-specific binding of MYC at promoter distal sites correlated with subtype-specific binding of Mediator 1 (MED1) (Fig. 1E), which is a key marker of enhancer activity (Whyte et al., 2013). In addition, signal transducer and activator of transcription 3 (STAT3) and ER, two key transcription factors in TNBC and ER+ breast cancer, respectively, co-occupied these subtype-specific MYC enhancers (Fig. 1E and Suppl. Fig. 1E). Given that MYC is not considered a pioneer factor (Soufi, Donahue, & Zaret, 2012), and it has been shown to accumulate in already active regions (Sabò et al., 2014), this suggests that subtype-specific enhancer-binding of MYC is primarily dictated by cancer type-specific driving transcription factors.

To investigate if MYC-bound enhancers are active within patients, we intersected them with assay for transposase-accessible chromatin with sequencing (ATAC-seq) peaks from TCGA breast cancer patients (Corces et al., 2018) (Fig. 1F). 72-77% of the MYC subtype-specific enhancers were identified in the TCGA patient ATAC-seq data. Importantly, TNBC- and ER+-specific MYC enhancers showed subtype-specific accessibility in patients indicating that they are indeed clinically relevant subtype-specific enhancers. This was further supported by subtype-selective expression of nearby genes in the TCGA patient expression data (Cancer Genome Atlas Research et al., 2013) (Suppl. Fig. 1F). Taken together, these findings suggest that oncogenic MYC invades cancer type-specific enhancers co-bound by cancer type-specific driving transcription factors.

### MYC enhancer target genes are cancer type-specific and prognostic

The findings above suggested that the reported cell type-specific role of MYC (Green et al., 2016; Ji et al., 2011) may be mediated by MYC enhancer binding rather than by its well-described role at promoters. To investigate this, we ranked putative MYC target genes based on the ratio of MYC binding at promoters compared to enhancers within 50kb (Fig. 2A). In addition to genes with the expected strong binding of MYC at their promoters, this analysis identified putative MYC target genes dominated by nearby MYC enhancer binding with no or minimal MYC binding at their promoters. This suggests that oncogenic MYC does not preferentially invade enhancers near its classical target promoters, but instead suggests that MYC binding at enhancers and promoters targets largely distinct gene programs. Enhancer-dominated target genes were also more highly connected to MYC enhancers through previously predicted enhancer-promoter pairs (Thurman et al., 2012) than promoter-dominated genes (Suppl. Fig. 2A), further implicating MYC enhancers in the regulation of these genes.

**Figure 2.**
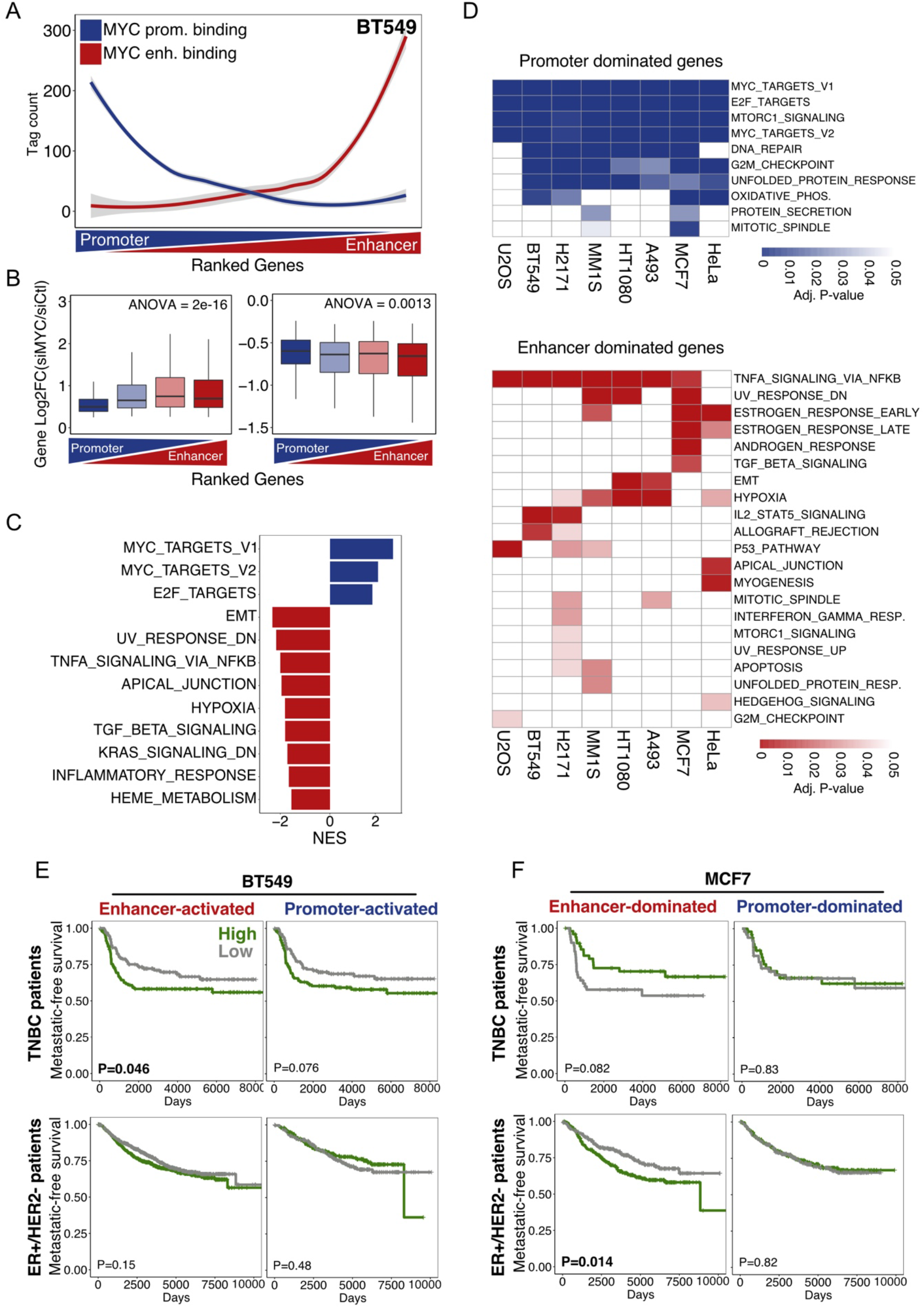
MYC enhancer binding controls cancer type-specific and prognostic gene programs. **(A)** Ranking of putative MYC target genes based on the enhancer/promoter ratio of MYC binding in BT549 cells. Enhancer binding of MYC was defined as the sum of binding at all enhancers within 50kb of the TSS. **(B)** RNA-seq was used to measure differential gene expression over the ranking of MYC promoter/enhancer associated genes from panel A in BT549 cells. The Log2FC for all significantly differentially expressed genes (adjusted P-value < 0.05) is shown for repressed and activated genes. The statistical difference in Log2FC was calculated with the ANOVA-test. **(C)** GSEA was performed over the promoter/enhancer ranking of differential expressed genes from RNA-seq as defined in panel B. Hallmark gene sets significantly enriched in enhancer- and promoter-dominated genes are represented in the bar plot by their Normalized Enrichment Score (NES). Blue (positive NES) represent MYC promoter-activated and red (negative NES) represent MYC enhancer-activated terms. All hallmarks presented are significantly enriched (adjusted P-value < 0.05). **(D)** The 25% most promoter- and enhancer-dominated genes were identified across 8 cancer cell lines from public MYC ChIP-seq data and GO-term enrichment analysis using Hallmark gene sets (Liberzon et al., 2015) was performed. Hallmarks gene sets that were significantly enriched (adjusted P-value < 0.05, hypergeometric test) within at least one cell line are presented. **(E)** Kaplan-Meier (KM) analyses using the Metabric cohort (Curtis et al., 2012) were performed for the top 25% MYC promoter- or enhancer-activated genes significantly repressed (adjusted P-value < 0.05 and Log2FC < -0.5) by MYC siRNA knockdown in BT549 cells. Mean expression of the gene sets was used to stratify ER+ and TNBC patients into high (top 50%) and low (bottom 50%) expression groups. Metastasis-free survival was plotted over days and the P-value was calculated using the log-rank test. **(F)** The same analysis as in panel E were performed on the top 10% MYC promoter- or enhancer-dominated genes. Mean expression of the gene sets was used to stratify ER+ and TNBC patients into high (top 20%) and low (bottom 20%) expression groups.

To assess the regulatory potential of MYC at enhancers, siRNA-mediated knockdown of MYC (Suppl. Fig. 2B) coupled with RNA-seq was performed. Differentially expressed genes were subdivided into four bins based on the relative binding of MYC to enhancers and promoters as described above (Fig. 2B). Interestingly, MYC enhancer-dominated genes were significantly more affected by MYC knockdown compared to promoter-dominated genes. This demonstrates that enhancer binding of MYC drives robust changes in the expression of nearby genes.

Gene set enrichment analysis (GSEA) of hallmark gene sets (Liberzon et al., 2015) in BT549 cells showed that MYC promoter-dominated genes were classical MYC targets, e.g., Cyclin-Dependent Kinase 6 (*CDK6*) and Cyclin-Dependent Kinase Inhibitor 1B (*CDKN1B*) (Z. Li et al., 2003), while enhancer-dominated genes were enriched for key features of TNBC such as Epithelial-Mesenchymal Transition (EMT) and hypoxia, e.g. C-X-C Motif Chemokine Ligand 8 (*CXCL8*) and Cadherin 3 (*CDH3*) (Yang, Zhao, Cui, & Liang, 2017; Yi, Peng, Xia, & Gan, 2022) (Fig. 2C). Similar results were observed in MCF7 cells, where MYC promoter-dominated genes were enriched for classical MYC targets, and MYC enhancer-dominated genes were enriched for estrogen-responsive genes characteristic of this breast cancer subtype (Suppl. Fig. 2C,D). To determine if this observation was a general trend across cancers, we performed Gene Ontology (GO) analyses of MYC promoter- and enhancer-dominated genes from six additional cancer cell lines (Fig. 2D). Consistent with our findings in breast cancer, MYC enhancer-dominated gene programs were indeed cancer type-specific across these cancer models. These findings suggest that MYC promoter binding drives the same classical MYC target gene program across cancers, whereas MYC enhancer binding drives distinct cancer type-specific gene expression.

Given the cancer type-specificity of MYC enhancer-dominated target genes, we speculated if these gene programs are clinically important in patients. To investigate this, we stratified patients from Metabric (Curtis et al., 2012) based on expression of enhancer- or promoter-dominated MYC target genes significantly activated by MYC in BT549 cells. Interestingly, high expression of MYC enhancer-activated genes in TNBC patients was significantly associated with poor prognosis, while MYC promoter-activated genes had slightly lower prognostic power (Fig. 2E). Importantly, these enhancer-activated genes from the TNBC BT549 cells were not associated with metastatic-free survival in ER+/HER2-breast cancer patients, indicating that MYC promotes a subtype-specific prognostic gene program from enhancers in TNBC. A similar analysis in luminal A MCF7 cells using all enhancer- and promoter-dominated genes showed that high expression of MYC enhancer-dominated genes predicted poor outcome specifically in ER+/HER2-breast cancer patients, but not in TNBC patients (Fig. 2F). In fact, high expression of these genes showed a tendency towards association with good outcome in TNBC. There was no association between expression of promoter-dominated MYC target genes and outcome for patients with ER+/HER2-tumors (Fig. 2F). These results demonstrate the clinical relevance of MYC enhancer binding and suggest that oncogenic MYC binds cancer type-specific enhancers to control cancer type-specific prognostic gene programs.

### MYC regulates enhancer activity together with cancer type-specific transcription factors

To study the direct transcriptional mechanisms of MYC at enhancers and avoid secondary effects of MYC perturbation (e.g., MYC-mediated effects on the cell cycle), we used a small molecule inhibitor, KJ-PYR-9, to acutely perturb MYC function. This inhibitor specifically blocks MYC-MAX heterodimerization, without affecting other related transcription factors (Hart et al., 2014). Given the profound effect of MYC on cancer cell transcription and proliferation, this acute perturbation of MYC function is critical to understand the direct effect of this oncoprotein at enhancers.

Three hours of treatment with KJ-PYR-9 effectively sequestered MYC completely away from both promoters and enhancers (Fig. 3A and Suppl. Fig. 3A). Importantly, acute gene regulation by KJ-PYR-9 was recapitulated by siRNA-mediated knockdown of MYC, and KJ-PYR-9 treatment acutely affected the expected gene programs similarly as more long-term siRNA-mediated knockdown of MYC (Suppl. Fig. 3B,C). MYC has previously been suggested to globally amplify RNA production (Nie et al., 2012), however, spike-in normalized RNA-seq illustrated that this was not an important mechanism in breast cancer cell lines using acute or more longterm siRNA-mediated MYC perturbation (Suppl. Fig. 3D). Taken together, this shows that we can use KJ-PYR-9 treatment to perturb MYC function acutely and specifically to directly address the role of this oncoprotein at enhancers.

**Figure 3.**
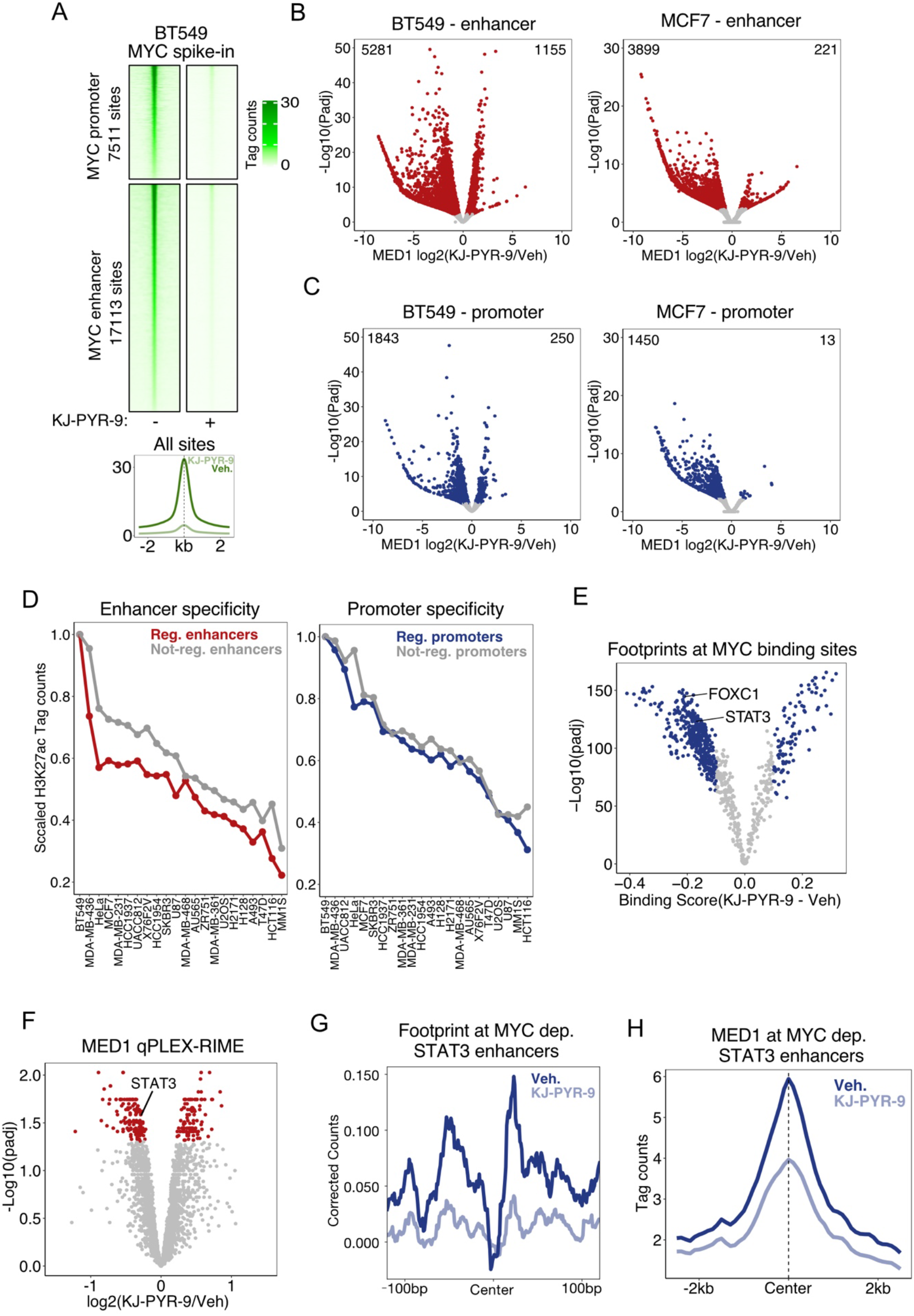
MYC modulates the activity of breast cancer subtype-specific enhancers. (A) Top, spike-in normalized MYC ChIP-seq in BT549 cells (n=3) treated with 40μM KJ-PYR-9 (3h) is visualized for MYC promoters and enhancers in a heatmap. Bottom, average line plot for all MYC binding sites. (**B-C**) MCF7 and BT549 cells were treated with KJ-PYR-9 (3h) and MED1 ChIP-seq was performed (n=3). Differential binding analyses of MED1 at MYC enhancer (**B**) and promoter **(C)** binding sites were visualized in volcano plots. Colored dots show significantly differential binding sites (adjusted P-value < 0.1, DiffBind). **(D)** The relative H3K27ac ChIP-seq signal across different cells lines (GSE65201 (Wang et al., 2015) and GSE96867 (Franco et al., 2018)) at enhancers and promoters occupied by MYC in BT549 cells is shown. MYC binding sites are subdivided into groups with significant and non-significant changes in MED1 binding in response to KJ-PYR-9 as determined in panel B and C. **(E)** Volcano plot showing changes in TOBIAS footprint scores (Bentsen et al., 2020) in response to KJ-PYR-9 treatment (3h) in BT549 cells. Significant footprints (adj. P-value < 10^−50^) are highlighted in blue. **(F)** MED1 qPLEX-RIME performed in BT549 cells treated with KJ-PYR-9 (3h) visualized in a volcano plot. Significantly changing proteins are highlighted in red (Padj<0.05, qPLEXanalyzer). **(G)** STAT3 footprint profiles for lost STAT3 footprints (Log2FC < -0.5) in MYC enhancers upon KJ-PYR-9 treatment (3h). **(H)** MED1 binding as determined by ChIP-seq at MYC enhancers with loss of STAT3 footprints upon KJ-PYR-9 treatment (see panel G).

To define MYC-regulated enhancers, we performed MED1 ChIP-seq in BT549 and MCF7 cells during acute 3h KJ-PYR-9 treatment. Differential binding analysis of MED1 at MYC binding sites demonstrated extensive loss of MED1 binding at these sites upon KJ-PYR-9 treatment with few sites showing a gain in MED1 binding (Fig. 3B,C). This suggests that MYC primarily promotes enhancer activation. Interestingly, MYC-activated enhancers had lower enrichment of E-boxes, but stronger MYC binding compared to unaffected MYC enhancers in BT549 cells (Suppl. Fig. 3E,F), indicating that direct MYC binding to E-boxes is not required for MYC function at enhancers. As expected from our analyses of all MYC-bound enhancers, MYC-regulated enhancers were highly cell type-specific between MCF7 and BT549 cells compared to promoters (Suppl. Fig. 3G). This high cancer type-specificity of MYC-regulated enhancers was further demonstrated using public H3K27ac ChIP-seq data from 12 cancer cell lines. Here, MYC-activated enhancers in TNBC BT549 cells were more cancer type-specific compared to unaffected MYC-bound enhancers (Fig. 3D). This increase in cancer type-specificity was not observed for MYC-regulated promoters (Fig. 3D). Taken together, these results demonstrate that MYC regulates enhancer activity in a highly cancer type-specific manner through a non-canonical mechanism less dependent on its classical binding motif. Consistent with our previous findings (Fig. 1E), a comprehensive Giggle analysis (Layer et al., 2018) identified breast cancer subtype-specific binding of transcription factors to MYC-activated enhancers (Suppl. Fig. 3H). This included YAP1, STAT3, and TEAD4 binding to MYC-activated enhancers in TNBC BT549 cells and ER, FOXA1, and GATA3 binding to MYC-activated enhancers in ER+ MCF7 cells.

To investigate if MYC regulates the binding of these subtype type-specific transcription factors to chromatin, we performed ATAC-seq in BT549 cells in response to KJ-PYR-9 treatment followed by footprinting analysis using the TOBIAS workflow (Bentsen et al., 2020). Consistent with the Giggle analysis, we found that MYC promotes the formation of footprints from previously described important transcription factors in TNBC such as STAT3 (Conway et al., 2020) and FOXC1 (Pan et al., 2018) transcription factors (Fig. 3E). Interestingly, MED1 qPLEX-RIME in response to KJ-PYR-9 treatment demonstrated that MYC inhibition abolished the association between STAT3 and MED1 (Fig. 3F). These findings suggest that MYC is required for STAT3 transcriptional function. Interestingly, the footprinting analyses demonstrated that STAT3 footprints were mostly lost at MYC-bound enhancers and not promoters (Suppl. Fig. 3I). Importantly, enhancers that lost STAT3 footprints in response to KJ-PYR-9 treatment also lost MED1 binding (Fig. 3G-H). Taken together, these findings indicate that MYC is required for binding and function of STAT3 at enhancers, which suggests that cooperativity between MYC and subtype-specific transcription factors is an important mechanism through which MYC controls cancer type-specific enhancer activity.

### Glucocorticoid hormone signaling directs oncogenic MYC enhancer activity

Enhancers are highly plastic and subject to change in response to extracellular stimuli from the tumor microenvironment (Shlyueva, Stampfel, & Stark, 2014). To determine if MYC activity at enhancers is also responsive to extracellular signals, we focused on glucocorticoid signaling. This has been shown to alter the enhancer landscape (McDowell et al., 2018) and to play an important role in TNBC by decreasing chemosensitivity (West et al., 2018) and promoting lung colonization (Obradović et al., 2019).

We first investigated the impact of the synthetic glucocorticoid Dexamethasone (Dex) on the MYC interactome by MYC qPLEX-RIME in BT549 cells (Fig. 4A). Interestingly, Dex treatment increased the association between the glucocorticoid receptor (GR) and MYC indicating a functional crosstalk between these transcription factors in response to extracellular glucocorticoids. In addition, MYC association with EST Variant Transcription Factor 6 (ETV6) and DExH-Box Helicase 30 (DHX30) was significantly changed, indicating that these factors may also play a role in shaping the Dex response together with MYC and GR.

**Figure 4.**
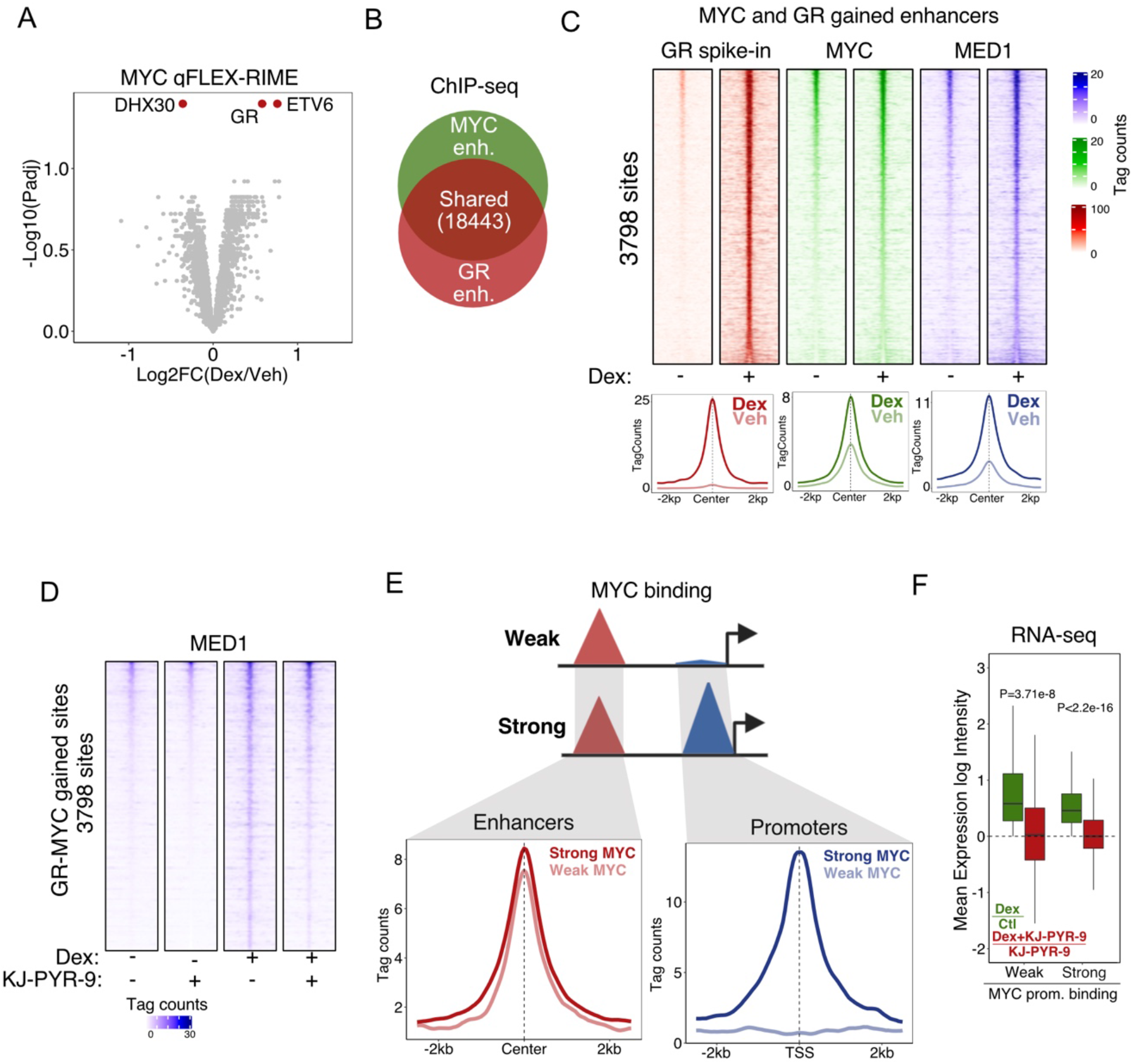
Glucocorticoid signaling directs MYC enhancer activity. **(A)** MYC qPLEX-RIME in BT549 cells in response to Dex treatment (2h) visualized in a volcano plot. Significantly changing proteins are indicated in red (Padj<0.05, qPLEXanalyzer). **(B)** Venn-diagram showing overlap between GR and MYC binding at enhancers in BT549 cells upon Dex treatment (2h) as determined by ChIP-seq. **(C)** Heatmap visualization (top) and average line plots (bottom) of GR, MYC, and MED1 ChIP-seq data at shared GR and MYC binding sites with a significant increase in MYC binding in response to Dex (2h) (adjusted P-value < 0.05, DESeq2). **(D)** Heatmap of MED1 ChIP-seq data in response to Dex (2h) and KJ-PYR-9 (3h) treatment (n=3) around the shared MYC and GR enhancers that gain MYC binding defined in panel C. **(E)** Genes within 50kb of shared MYC-GR enhancers were subdivided depending on the binding of MYC at their promoter (high=1946 sites and low=760 sites). The average plots (bottom) display MYC binding at enhancers and promoters of the two groups as determined by ChIP-seq. **(F)** RNA-seq was performed in response to Dex (2h) and KJ-PYR-9 (3h) to evaluate the effect on the gene groups defined in panel E. P-values were calculated using Welch’s t-test, two-sided.

To further explore the crosstalk between GR and MYC at enhancers, we performed ChIP-seq with specific antibodies against GR, MYC, and MED1 in BT549 cells treated with Dex. Consistent with the qPLEX-RIME data, we identified extensive overlap in the binding of GR and MYC at gene-distal regions in response to Dex treatment (Fig. 4B and Suppl. Fig. 4A). These regions were highly occupied by MED1 (Suppl. Fig. 4A), indicating that these gene-distal shared GR and MYC regions were mostly active enhancers. Interestingly, although Dex increased MYC expression (Suppl Fig. 4B), we only found a significant increase in MYC binding in response to Dex-treatment at a small subset (3,798) of shared GR-MYC enhancers, i.e., gained GR-MYC enhancers (Fig. 4C). This indicates that GR specifically directs MYC binding to these enhancers. These gained GR-MYC enhancers showed a dramatic Dex-mediated increase in enhancer activity as determined by MED1 binding (Fig. 4C), indicating that these enhancers play a critical role in the transcriptional response to Dex. Interestingly, KJ-PYR-9 treatment abrogated the Dex-mediated induction of MED1 binding at these enhancers demonstrating that MYC is required for GR-mediated activation of enhancer activity (Fig. 4D). These findings demonstrate that GR can direct MYC binding to new enhancers, where MYC cooperates with GR to activate these enhancers.

To more directly investigate if cooperation between MYC and GR at enhancers is also required to activate the GR target gene program, genes within 50kb of gained GR-MYC enhancers were subdivided into two groups depending on their MYC binding intensity at the promoter (Fig. 4E, top). This provided two gene groups with (strong) and without (weak) MYC promoter binding, but with similarly strong gained GR-MYC enhancers nearby (Fig. 4E, bottom). Interestingly, activation of Dex-induced genes in both groups was highly impaired by co-treatment with KJ-PYR-9 as determined by RNA-seq (Fig. 4F). This demonstrates that cooperativity between GR and MYC at enhancers plays a critical role for the transcriptional response to Dex. Interestingly, many of the target genes of gained GR-MYC enhancers are involved in metastatic features of TNBC e.g., *CXCL8*, Fibroblast Growth Factor 2 (*FGF2*), and Zinc Finger E-Box Binding Homeobox 1 (*ZEB1*). This indicates that GR-MYC cooperativity at shared enhancers in response to extracellular glucocorticoids induces aggressive cancer features in TNBC.

### MYC modulates RNAPII recruitment and eRNA production at enhancers

It is well-established that MYC can regulate pause-release at target gene promoters (Rahl et al., 2010). Interestingly, it has recently been shown that pause-release also plays an important role in enhancer transcription (Henriques et al., 2018). Thus, we hypothesized that MYC controls enhancer activity by regulating this process. To investigate this, we performed ChIP-seq in response to KJ-PYR-9 treatment for RNAPII phosphorylated on serine 5 (S5p) and serine 2 (S2p), which mark initiating/paused and elongating RNAPII, respectively. We focused our analyses on comparing the previously described MYC-activated enhancers (Fig. 3B) and MYC-bound promoters where downstream gene expression was significantly decreased by KJ-PYR-9. Consistent with previous findings (Rahl et al., 2010), we found that MYC is important for efficient pause-release of RNAPII at MYC-bound promoters of MYC-activated genes (Fig. 5A,B). This was not a global effect of KJ-PYR-9, but specific to MYC-bound and regulated genes (Suppl. Fig. 5A). In contrast, KJ-PYR-9 induced loss of both phosphorylation states of RNAPII at enhancers, and the S5p/S2p RNAPII ratio was decreased in response to KJ-PYR-9 indicating less pausing upon MYC inhibition (Fig. 5A,B). Taken together, this suggests that MYC is important for RNAPII recruitment at enhancers rather than promoting pause-release as observed at promoters. This is further supported by analysis of published data from U2OS and P493-6 cells (Lin et al., 2012; Walz et al., 2014) showing increased RNAPII binding at MYC enhancers upon MYC overexpression (Fig. 5C).

**Figure 5.**
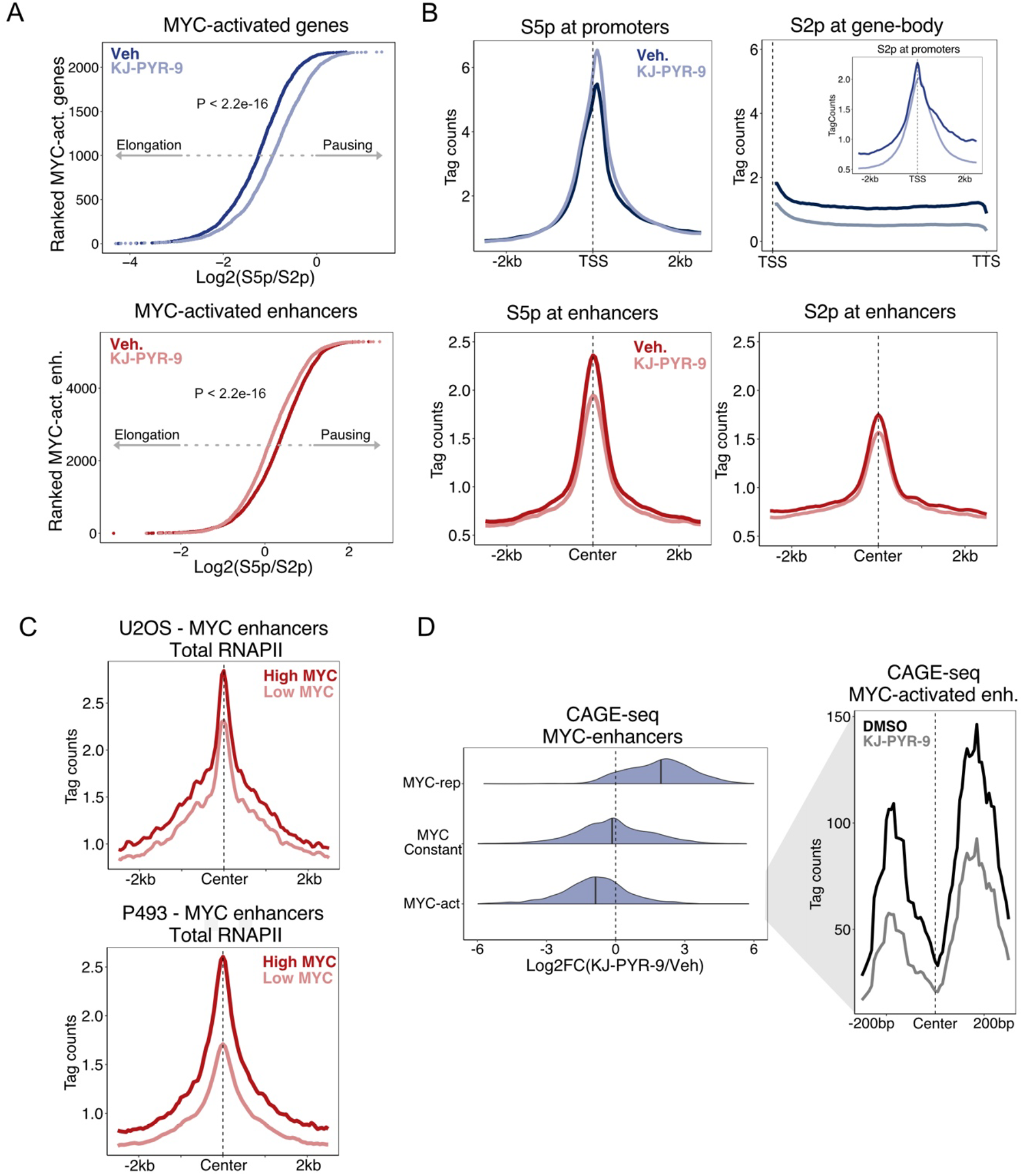
MYC controls RNAPII recruitment rather than pause-release at enhancers to promote enhancer RNA expression. **(A)** MYC-activated genes were defined as genes significantly repressed by KJ-PYR-9 treatment (3h) as determined by RNA-seq (Padj<0.05 and log2FC>0, DESeq2) with MYC promoter binding as determined by ChIP-seq. MYC-activated enhancers were defined as promoter-distal MYC binding sites that gain MED1 binding in response to KJ-PYR-9 (3h) as determined by ChIP-seq (Padj<0.05, DiffBind, see Fig. 3B). The MYC-activated promoters and enhancers were then ranked based on their pausing ratios defined by the RNAPII S5p/S2p ratio as determined by ChIP-seq in BT549 cells in response to KJ-PYR-9. The S5p ChIP-seq signal was counted +/-1kb of the TSS and the S2p signal was counted 10kb into the gene body from promoters. For enhancers, both S5p and S2p were counted +/- 500bp around the center of the MYC binding sites within the enhancer regions. P-values were calculated using Welch’s t-test, two-sided. **(B)** Average plots show the RNAPII S5p and S2p occupancy as determined by ChIP-seq around the MYC-activated genes with MYC promoter binding and MYC-activated enhancers (see Fig. 3B). **(C)** Total RNAPII ChIP-seq in enhancers that become occupied by MYC upon overexpression in U2OS and P493-6 cells (GSE44672 (Walz et al., 2014) and GSE36354 (Lin et al., 2012)). **(D)** CAGE-seq in response to KJ-PYR-9 treatment (3h) was performed. Left, Log2FC are plotted for MYC-activated, consistent and MYC-repressed enhancers measured by change in MED1 binding (see Fig. 3B). Right, zoom in on the average CAGE-seq profile around the MYC-activated enhancers.

It is well-established that RNAPII binding at enhancers results in the production of small non-coding RNAs termed enhancer RNA (eRNA), which is a hallmark of active enhancers (De Santa et al., 2010; T. K. Kim et al., 2010). To investigate if the decreased recruitment of RNAPII in response to MYC inhibition also leads to a decrease in eRNA production from these enhancers, we performed Cap Analysis Gene Expression (CAGE-seq) to map the 5’-ends of nascent RNA (Shiraki et al., 2003). Interestingly, MYC-activated enhancers as determined by MED1 binding showed decreased eRNA production in response to KJ-PYR-9 (Fig. 5D). This was supported by Ribo-Zero RNA-seq showing decreased eRNA levels in response to KJ-PYR-9 at these enhancers (Suppl. Fig. 5B). This decrease in eRNA production was associated with a decrease in chromatin accessibility (Suppl. Fig. 5C), consistent with previous findings at selected enhancers (Mousavi et al., 2013). Altogether, this demonstrates that MYC plays a key role in controlling enhancer transcription by inducing recruitment of RNAPII, which is different from its role in promoting pause-release at promoters.

### MYC regulates an H3K9 epigenetic switch to promote enhancer activity

To further unravel the role of MYC at enhancers and understand how MYC induces RNAPII recruitment and eRNA production, we re-visited the MED1 qPLEX-RIME (Fig. 3F). One interesting candidate from this experiment was KDM3A, which showed decreased association with MED1 upon KJ-PYR-9 treatment. KDM3A is a lysine demethylase known to target lysine 9 on histone 3 (Wilson, Fan, Sahgal, Qi, & Filipp, 2017), and we speculated that this epigenetic modifier could play a role in MYC-mediated transcriptional regulation. KJ-PYR-9 treatment led to a global loss of KDM3A from chromatin as determined by ChIP-seq, particularly at MYC-bound enhancers (Fig. 6A and Suppl. Fig. 6A). This was consistent with a modest decrease in KDM3A protein expression in response to MYC inhibition (Suppl. Fig. 6B). Interestingly, despite a global loss of KDM3A from chromatin in response to KJ-PYR-9, its histone substrates, H3K9me1 and H3K9me2, were specifically increased at MYC-activated enhancers and not at MYC-activated genes (Fig. 6B). This suggests that KDM3A plays an important role in demethylating H3K9me1/2 specifically at enhancer regions, despite its more robust binding at promoters (Suppl. Fig. 6C).

**Figure 6.**
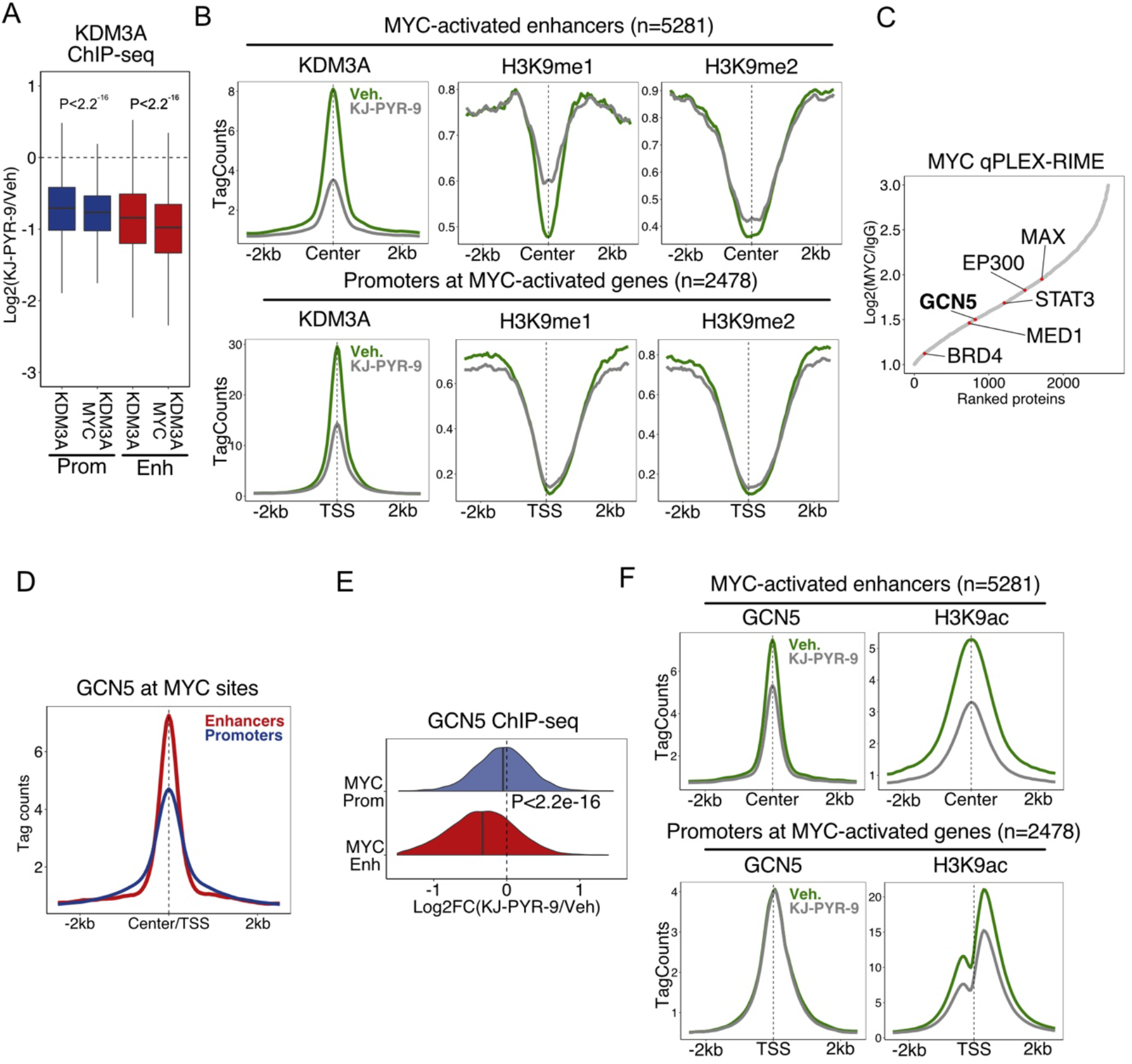
MYC activates an H3K9 epigenetic switch at enhancers. **(A)** Boxplot showing the Log2FC in KDM3A binding with KJ-PYR-9 treatment (3h) as determined by ChIP-seq in BT549 cells at shared (KDM3A and MYC) or KDM3A-only enhancers and promoters. P-values were calculated using Welch’s t-test, two-sided. **(B)** Average line plots showing KDM3A, H3K9me1/me2 occupancy as determined by ChIP-seq at the MYC-activated enhancers and MYC-activated genes (see Fig. 5A and Fig. 3B). **(C)** Proteins identified as MYC-associated proteins by qPLEX-RIME (same data as Fig. 4A) were ranked according to their fold enrichment over IgG control. Selected proteins are highlighted. **(D)** ChIP-seq binding intensity of GCN5 at MYC-bound promoters and enhancers in BT549 cells as determined by ChIP-seq. (**(F)** Change in GCN5 binding in response to KJ-PYR-9 treatment (3h) at MYC promoters and enhancers. P-value was calculated using Welch’s two-sided t-test. (**E**) Average line plots showing GCN5, H3K9ac occupancy as determined by ChIP-seq at MYC-activated enhancers and MYC-activated genes (see Fig. 5A and Fig. 3B).

Surprisingly, we did not identify KDM3A in our MYC qPLEX-RIME experiment indicating that KDM3A is not part of the MYC complex. Interestingly, however, we robustly identified the acetyltransferase GCN5 in our MYC qPLEX-RIME (Fig. 6C), which has previously been shown to be a critical and highly specific regulator of H3K9ac (Jin et al., 2011). Thus, we speculated that MYC recruits GCN5 to enhancers, where it, together with KDM3A, drives an epigenetic switch to a permissive chromatin structure characterized by low H3K9me1/2 and high H3K9ac.

To investigate if MYC directly recruits GCN5 to enhancers, we performed ChIP-seq in response to KJ-PYR-9 treatment. 83% of the identified GCN5 binding sites overlapped with MYC binding, with the strongest signal found at MYC enhancers (Fig. 6D and Suppl. Fig. 6D). Interestingly, without changing GCN5 protein levels (Suppl. Fig. 6B), KJ-PYR-9 treatment led to a specific loss of GCN5 binding at MYC enhancers and no change in binding at promoters (Fig. 6E). This suggests that MYC specifically recruits GCN5 to enhancer regions. Importantly, loss of GCN5 recruitment at MYC-activated enhancers was associated with a strong decrease in H3K9ac (Fig. 6F). Taken together, these findings suggest that MYC induces an epigenetic switch at H3K9 around target enhancers through the combined action of KDM3A and GCN5.

### MYC-induced epigenetic switch at enhancers promotes BRD4 binding and BRD4-mediated RNAPII recruitment

We speculated that MYC-induced H3K9ac at target enhancers may lead to the recruitment of BRD4, which is known to bind acetylated histones, including H3K9ac (Devaiah et al., 2016), and acetylated transcription factors through its bromodomains (Dey, Chitsaz, Abbasi, Misteli, & Ozato, 2003). In support of this, we identified BRD4 in our MYC qPLEX-RIME experiment (Fig. 6C). To investigate if MYC promotes BRD4 recruitment at enhancers, we performed BRD4 ChIP-seq in response to KJ-PYR-9 treatment (Fig. 7A). This demonstrated that MYC specifically recruits BRD4 to enhancers and not promoters without altering BRD4 expression (Suppl. Fig. 7A).

**Figure 7.**
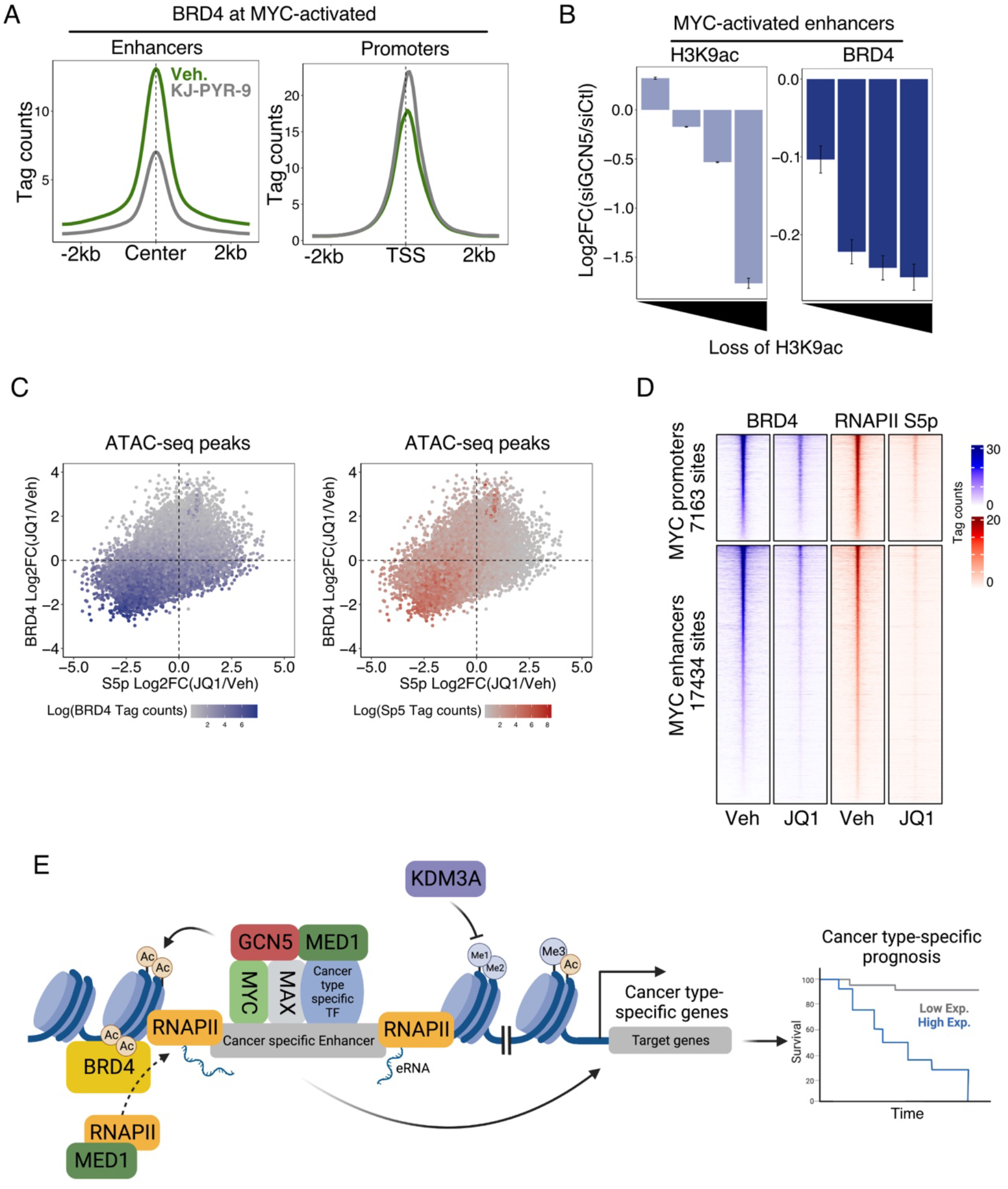
MYC promotes RNAPII recruitment to enhancers through a GCN5-BRD4-axis. **(A)** BRD4 ChIP-seq signal at MYC-activated enhancers and promoters in response to KJ-PYR-9 treatment (3h) in BT549 cells. **(B)** MYC-activated enhancers were divided into quartiles depending on the Log2FC of H3K9ac in response to siRNA-mediated knock down of GCN5 as determined by ChIP-seq. The change in BRD4 recruitment based on ChIP-seq was plotted in the same bins. Error bars represent SEM. **(C)** Scatterplot showing the Log2FC of BRD4 and RNAPII S5p ChIP-seq signal upon JQ1 treatment (1h) in BT549 cells within ATAC-seq peaks (n=46010). The dot color represents the Tag counts intensity of BRD4 and RNAPII S5p binding at the given site. **(D)** Heatmap of BRD4 and RNAPII S5p binding as determined by ChIP-seq in response to JQ1 treatment (1h) in BT549 cells within MYC-bound enhancers and promoters. **(E)** Model of MYC function at enhancers. MYC binds cancer type-specific enhancers together with cancer type-specific transcription factors to regulate prognostic gene programs important for the specific cancer phenotype. MYC recruits GCN5 to acetylate H3K9, which promotes recruitment of BRD4 specifically to MYC enhancers. BRD4 together with MED1 binding mediates the recruitment of RNAPII leading to increased eRNA transcription and thereby enhancer activity.

To investigate if GCN5-induced acetylation was important for BRD4 recruitment, we performed H3K9ac and BRD4 ChIP-seq upon siRNA-mediated knock down of GCN5. Importantly, GCN5 knockdown did not alter the levels of MYC, H3K9ac, or BRD4 (Suppl. Fig. 7B). Interestingly, loss of H3K9ac upon GCN5 depletion was associated with loss of BRD4 recruitment at MYC enhancers (Fig. 7B). Thus, although other mechanisms (e.g., other acetylated histone marks) are likely also involved in recruiting BRD4, these findings indicate that MYC-mediated recruitment of GCN5 and subsequent H3K9ac plays an important role in recruiting BRD4 to MYC-activated enhancers.

BRD4 has been shown to associate strongly with active transcriptional sites and mediate the recruitment of RNAPII through its interaction with the mediator complex (Bhagwat et al., 2016; S.-Y. Wu & Chiang, 2007). To investigate if BRD4 is also involved in the recruitment of RNAPII to MYC-bound enhancers, we performed BRD4 and RNAPII S5p ChIP-seq upon acute 1h JQ1 treatment, which blocks the BRD4 bromodomains that bind to acetylated residues (Filippakopoulos et al., 2010). Comparing BRD4 and RNAPII S5p recruitment to all accessible chromatin sites in BT549 cells demonstrated that strong BRD4 binding sites were lost upon JQ1 treatment, and this resulted in loss of RNAPII S5p occupancy (Fig. 7C). Consistent with this global analysis, specifically focusing on MYC-bound regions showed that blocking the bromodomains of BRD4 with JQ1 led to a dramatic loss of BRD4 and RNAPII recruitment to both MYC-bound promoters and enhancers (Fig. 7D) without changing the global levels of BRD4, GCN5, MYC, or H3K9ac (Suppl. Fig. 7C). Taken together, these results demonstrate that activation of cancer type-specific enhancers by MYC involves an epigenetic switch that promotes BRD4-mediated RNAPII recruitment (Fig. 7E).

## Discussion

Oncogenic MYC has previously been shown to invade enhancer regions (Lin et al., 2012; Sabò et al., 2014; See et al., 2022; Walz et al., 2014; Zeid et al., 2018), but this MYC enhancer invasion has been suggested to be spillover from saturated MYC binding at promoters with low or no gene-regulatory potential (Nie et al., 2012; Walz et al., 2014). In contrast to this model, we demonstrate that MYC enhancer binding strongly activates oncogenic gene programs in a highly cancer type-specific manner. This is in line with previous findings for MYCN in neuroblastoma (Zeid et al., 2018). We show that key oncogenic transcription factors can direct MYC enhancer binding, which is consistent with recent findings showing that MYC and its paralogs depend on other transcription factors to access chromatin, e.g., TWIST1 (Zeid et al., 2018), WDR5 (Thomas et al., 2019). In turn, we show that MYC supports the binding and function of the oncogenic transcription factors STAT3 and GR in TNBC. Based on these findings, we propose that MYC supports key oncogenic transcription factors to promote activation of cancer type-specific enhancers driving cancer type-specific gene programs. Importantly, these MYC enhancer target gene programs are also predictive of patient outcomes in a cancer type-specific manner, indicating that MYC enhancer invasion plays a clinically important role in regulating tumor progression. Thus, we propose a revised gene-specific affinity model (Lorenzin et al., 2016), where MYC target gene specificity is largely determined by cooperativity with lineage-determining transcription factors at enhancers and is less dependent on MYC-binding to promoters.

MYC is known to play an important role in regulating pause-release at target promoters by recruiting the P-TEFb complex containing CDK9 that catalyzes RNAPII serine 2 phosphorylation (Rahl et al., 2010). Recently, it was shown that P-TEFb-mediated pause-release is also an important mechanism regulating enhancer transcription (Henriques et al., 2018), which is tightly linked to enhancer activity (T. K. Kim et al., 2010). Surprisingly, we show that MYC induces transcription of eRNA by recruiting RNAPII rather than by increasing RNAPII pause-release as it does at promoters. This is driven by an enhancer-specific epigenetic switch from H3K9me1/2 to H3K9ac that promotes BRD4 binding and BRD4-mediated RNAPII recruitment. These findings demonstrate that MYC controls transcription from enhancers and promoters through different mechanisms. The reason for this difference between MYC function at enhancers and promoters is currently unclear. The non-canonical mechanism of MYC recruitment to enhancers, which is less dependent on E-boxes compared to binding of MYC at promoters, may play a role. Alternatively, differences in the chromatin-associated proteins between enhancers and promoters dictated by the different composition of DNA motifs at these regions (Andersson et al., 2014; Andersson & Sandelin, 2020) could affect MYC function. This is consistent with the recently proposed coalition model suggesting that the MYC interactome is key determinant of MYC function (Lourenco et al., 2021). In any case, MYC-mediated recruitment of MED1/BRD4/RNAPII and subsequent eRNA transcription is likely to play an important role in regulating target genes of MYC enhancer invasion. Transcription at enhancers has been shown to increase chromatin accessibility (Mousavi et al., 2013), which is consistent with our finding that loss of RNAPII recruitment in response to MYC inhibition is associated with decreased chromatin accessibility. In addition, many roles for eRNAs have been proposed, including stabilization of weak protein-DNA interactions to increase the transcriptional output and promotion of chromatin loops (Arnold, Wells, & Li, 2020). Consistently, MYC overexpression has recently been shown to regulate chromatin loop formation (See et al., 2022). Furthermore, it is particularly interesting to note that eRNAs have recently been proposed to stimulate RNAPII elongation by releasing Negative Elongation Factor (NELF) at target gene promoters (Gorbovytska et al., 2022). This indicates potential crosstalk between MYC at enhancers and promoters, where MYC-induced eRNA regulates pause-release at target gene promoters.

In conclusion, we show that MYC activates cancer type-specific enhancers driving prognostic gene programs by inducing an enhancer-specific epigenetic switch that promotes BRD4-mediated RNAPII recruitment and eRNA transcription.

## Methods

### Cell culture of breast cancer cell lines

MCF7, T47D, MDA-MB-231, and BT549 (ATCC) were grown in RPMI (Sigma, R8758) supplemented with 10% fetal bovine serum (Sigma-Aldrich, F7524), 50U/mL penicillin, 50μg/mL streptomycin (Lonza, DE17-602E) and 2mM L-glutamine. Cells were genotyped by short-tandem repeat genetic profiling (STR) using the PowerPlex_16HS_Cell Line panel and routinely mycoplasma tested. All cells were cultured at 37°C and 5% CO_2_.

### siRNA-mediated knockdown

BT549 cells were transfected with ON-TARGETplus SMARTPools (Dharmacon) targeting c-MYC (L-003282-02), GCN5 (L-009722-02) or non-targeting control (D-001810-01) using Lipofectamine RNAiMAX (Invitrogen, 13778100) at a final concentration of 10nM. The media was replaced 24h after transfection and the cells were harvested 48h later.

### MYC:MAX inhibition with KJ-PYR-9

MCF7 and BT549 cells were treated with 40 μM KJ-PYR-9 (MedChemExpress, HY-19735) dissolved in DMSO (Sigma, D8418) for 3h before harvest.

### BRD4 bromodomain inhibition with JQ1

BT549 cells were treated with 1μM JQ1 (MedChemExpress, HY-13030) dissolved in DMSO for 1h before harvest.

### GR activation with dexamethasone

BT549 cells were treated with 0.5μM Dexamethasone (MedChemExpress, HY-14648), dissolved in water, for 2h before harvest.

### Western blotting

Cells were washed twice in cold PBS and harvested in lysis buffer (50mM Tri-HCL pH 6.8, 10% Glycerol, 2.5% SDS, 10mM DTE, 10mM beta-glycerophosphate, 10mM NAF, 0.1mM Sodium Ortovandatate). The lysate was benzonase (Millipore, 70664) treated for 30 min. 20 μg protein was loaded in each lane on a 7% or 10% SDS-PAGE gel. Proteins were transferred to a nitrocellulose membrane, which was blocked in 5% skim milk in TBS with 0.1% Tween-20 and incubated with primary antibody overnight at 4°C. After washing the membrane, the secondary antibody was added for 1h before the membrane was washed and proteins detected using ECL. Primary antibodies: KDM3A (Bethyl Laboratories, A301-538A, 1:1000), MYC (Cell Signaling, 13987, 1:1000), β-tubulin (Sigma Aldrich, 05-661, 1:20000), GCN5 (Abcam, Ab217876, 1:1000), H3 (Cell Signaling, 2650, 1:2500), H3K9ac (Abcam, Ab4441, 1:1000), H3K9me1 (Abcam, Ab9045, 1:1000), H3K9me2 (Abcam, Ab176882, 1:1000), BRD4 (Bethyl Laboratories, A301-985A100).

### Ribosome depleted RNA-seq

Total RNA was harvested from cells with Tri-reagent (Sigma-Aldrich, 93289) and purified with chloroform on EconoSpin Columns (EconoSpin, 1910). Ribosomal RNA was depleted using the NEBNext rRNA depletion kit (Biolabs, E6310) before the library was prepared with the NEBNext RNA Library Prep kit (Biolabs, E7540). Samples were sequenced on a NovaSeq 6000 (Illumina) to reach approximately 25M paired-end reads per sample.

50bp paired-end reads were aligned to the human genome version GRCh38 using Hisat2 version 2.1.0. (D. Kim, Langmead, & Salzberg, 2015). Aligned reads were counted with feature-Counts version 1.6.4. (Liao, Smyth, & Shi, 2014) and differentially expressed genes were identified based on the negative binomial distribution using DESeq2 (Love, Huber, & Anders, 2014) in R.

### qPLEX-RIME

qPLEX-RIME was performed as described previously (Papachristou et al., 2018) with five biological replicates using MED1 (Bethyl Laboratories, A300-793A) or GR (Cell Signaling, D6H2L) antibodies. In brief, for each chromatin immunoprecipitation cells of two 15cm plates were crosslinked at RT using 2mM disuccinimidyl glutarate (DSG) for 20min followed by 1% formaldehyde for 10min, which was quenched in 0.1M glycine for 10min. Cells were washed twice in cold PBS and scraped off the plates in cold PBS containing protease inhibitors (Roche, Complete 11836170001 and Halt, Thermo Scientific, 78438). Crosslinked material was resuspended in lysis buffer 1 (50mM Hepes-KOH pH 7.5, 140mM NaCl, 1mM EDTA, 10% Glycerol, 0.5% NP-40, 0.25% Triton X-100) for 10min followed by 5min in lysis buffer 2 (10mM Tris-HCL pH 8.0, 200mM NaCl, 1mM EDTA, 0.5mM EGTA) before resuspending in lysis buffer 3 (10mM Tris-HCL pH 8.0, 100mM NaCl, 1mM EDTA, 0.5mM EGTA, 0.1% Na-Deoxycholate, 0,5% N-lauroylsarcosine). Samples were sonicated using Covaris Ultrasonicator ME220 until DNA fragments had a size between 150-500bp. Chromatin was immuno-precipitated overnight using protein A coated dynabeads (Dynabeads, 10002D) with specific antibodies. Beads were washed 10 times in cold RIPA buffer (50mM HEPES pH 7.6, 1mM EDTA, 0.7% Na-deoxycholate, 1% NP-40, 0.5M LiCl) followed by two washes in AMBIC buffer (100mM ammonium hydrogen carbonate). The bound protein complexes were digested on beads overnight using trypsin at a final concentration of 15ng/μL (Pierce, 90057) followed by a second digestion step the next day for 4h. Peptides were cleaned and labeled with the TMT-11plex reagents (Thermo Fisher Scientific), followed by fractionation using Reverse-Phase spin columns at high pH. LC-MS analysis was performed with an Easy nanoLC system (Thermo Fisher Scientific) coupled in line to a Q Exactive HF orbitrap instrument (Thermo Fisher Scientific). MS scan range was set to 400-1600m/z and the resolution was 120000 (MS). The top 20 peptides were subjected to MS^2^ with a resolution of 60000, isolation windows of 1.2m/z, a dynamic exclusion for 40s, and accepted charge states 2-6. Data was processed in Proteome Discoverer 2.5 (Thermo Fisher Scientific), using the search engine Mascot 2.7. The analysis included the following parameters: Precursor mass tolerance: 10ppm, fragment mass tolerance: 0.03Da, trypsin: 2 missed cleavages, static modification: TMT, dynamic modification: oxidation (M), deamidation (NQ), integration tolerance: 20ppm, SwissProt database for homo sapiens and Quan values correction was applied. qPLEX-RIME data was further analyzed using the qPLEXanalyzer workflow (Papachristou et al., 2018) in R using a limma-based analysis (Ritchie et al., 2015) to identify significant changes in protein interactions. Results were visualized using ggplot2 (Wickham, 2016).

### ChIP-seq

ChIP-seq samples were prepared in 2-3 biological replicates. Chromatin was prepared as described above for qPLEX-RIME. Immunoprecipitations were performed using specific antibodies for MED1, MYC, POL II Ser2p (Cell Signaling, 13499), POL II Ser5p (Cell Signaling, 13523), KDM3A, GCN5 (Abcam, Ab217876), BRD4, H3K9ac (Abcam, Ab4441), H3K9me1 (Abcam, Ab9045), H3K9me2 (Abcam, Ab176882). In indicated experiments, spike-in chromatin was used according to manufactories protocol (Activemotif, 61686 and 53083). After washing the beads, chromatin was eluted and decrosslinked by incubating at 65°C overnight in elution buffer (50mM Tris-HCL, 10mM EDTA, 1% SDS). Samples were treated with RNase A (20ng/mL) for 1h followed by proteinase K (200ng/mL) for 2h before DNA was purified by standard phenol-chloroform purification. Purified DNA was submitted to library preparation using NEBNext Ultra II DNA Library Prep Kit (New England BioLabs, E765). The samples were sequenced on a NovaSeq 6000 (Illumina) to reach approximately 25M paired-end reads per sample.

50bp paired-end reads were aligned to the human genome using Hisat2 version 2.1.0. PCR duplicated reads were removed using Samtools (Li et al., 2009) before peak calling was performed with the default setting in MACS2 (Zhang et al., 2008). Only peaks called within all replicates were considered for downstream analysis. Diffbind (Ross-Innes et al., 2012) was used for differential peak calling. Homer (Heinz et al., 2010) was used to create heatmaps, average plots, and for counting reads within regions. Read density is represented as Tag Counts, which is normalized to 10 million mapped reads.

### ATAC-seq

50,000 cells were lysed in ATAC-RSB (resuspension buffer) (10mM Tris-HCL pH 7.4, 10mM NaCl, 3mM MgCl_2_, 0.1% Tween-20) supplemented with 0.1% IgePal-CA630 and 0.01% Digitonin before being washed three times in cold ATAC-RSB buffer. Nuclei were pelleted and resuspended in 50μL transposition mixture (17.75μL PBS, 5uL Nuclease Free Water, 25uL TD 2x reaction buffer, 0.5uL 1% digitonin, 0.5uL 10% Tween-20, 1.25μL TDE1 (Illumina, 20034197)) and incubated at 37°C in a thermomixer at 1000RPM for 30min. DNA was cleaned using Qiagen PCR Purification kit before PCR amplification for 10 cycles using Illumina index primers, 25μL Q5 High Fidelity 2x master mix (New England BioLabs, M0492) in a final volume of 50μL. Samples were size selected and purified using Ampure XP beads (Bechman Coulter, A63881). Next-generation sequencing was performed using Illumina Novaseq 6000 to get ∽25M 50bp paired-end reads per sample. The analysis was performed similar to ChIP-seq described above, however, duplicate reads were included in the analysis. In general, all analyses were made using the Homer toolbox and visualized using ggplot2.

### CAGE-seq

BT549 cells treated with 40μM KJ-PYR-9 for 3h were subject to CAGE-seq analysis performed following a previously described protocol (Cvetesic et al., 2018). In brief, sample RNA was mixed with carrier RNA followed by first-strand cDNA synthesis. The ends of the RNA were oxidized, biotinylated and RNA fragments not overlapping first-strand cDNA were RNase digested. This provided a library of fragments where the biotinylated 5’cap was pulled down using streptavidin paramagnetic beads. The cap-trapped cDNA strand was then released by RNase treatment and the linker was ligated to the first cDNA strand followed by second-strand cDNA synthesis. The carrier was degraded, and the libraries were then purified using SPRI paramagnetic beads, PCR amplified and subjected to sequencing on Illumina NextSeq 550 to a minimum of 35M reads per sample. Analysis of the data was performed within the CageFigthR software (Thodberg, Thieffry, Vitting-Seerup, Andersson, & Sandelin, 2019)

## Acknowledgement

The authors thank Tenna Pavia Mortensen, Maibrith Wishoff, and Ronni Nielsen at the Functional Genomics and Metabolism research unit at SDU for expert technical support with Illumina sequencing. Work in the R.S. laboratory was supported by the Novo Nordisk Foundation (NNF18OC0053276 and NNF21OC0071373) and the Danish Cancer Society (R231-A13786). The mass spectrometry and proteomics platform at SDU, used to perform qPLEX-RIME, is financed by grants from the VILLUM Foundation (VILLUM Center for Bioanalytical Sciences, grant no. 7292 to O.N.J.), the Novo Nordisk Foundation (INTEGRA grant no. NNF20OC0061575 to O.N.J), and by PRO-MS: Danish National Mass Spectrometry Platform for Functional Proteomics (grant no. 5072-00007B to O.N.J). The work in the R.A. laboratory was supported by Novo Nordisk Foundation (NNF20OC0059796).

---

**Supplementary Figure 1.**
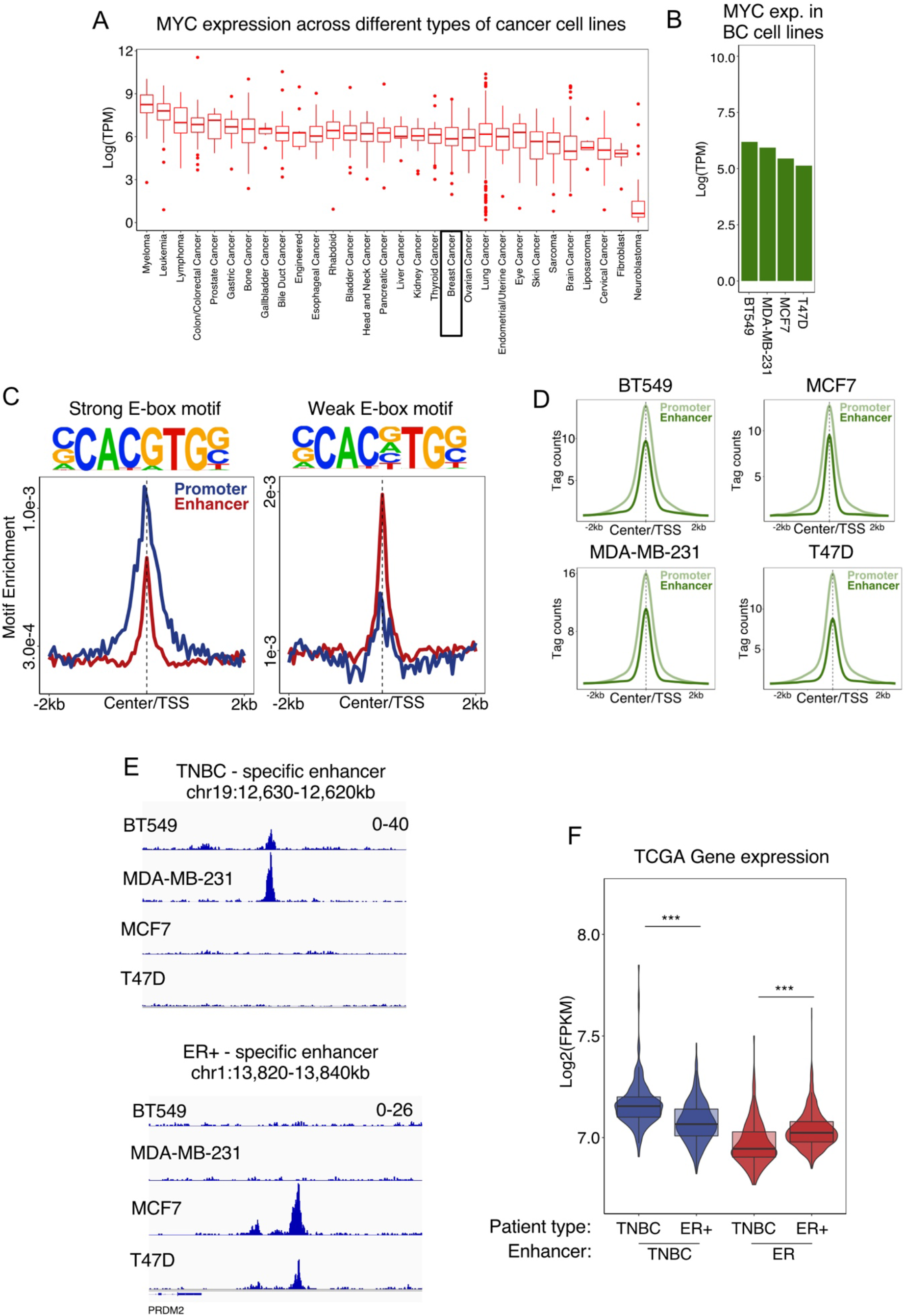
**(A)** MYC expression across cancer cell lines representing different tumor types from the DepMap database (Barretina et al., 2012) was summarized and plotted. The plot illustrates Log2 TPM counts. **(B)** MYC expression Log2(TPM counts) in BT549, MDA-MB-231, MCF7, and T47D cell lines. Data from the DepMap database. **(C)** Occurrence of strong and weak E-box motifs within MYC-bound promoters and enhancers from regions defined in Fig. 1A. **(D)** Average line plot of MYC ChIP-seq signal at MYC-bound promoters and enhancers across the four breast cancer cell lines BT549, MDA-MB-231, MCF7, and T47D. **(E)** Example of MYC breast cancer subtype-selective enhancers defined in Fig. 1E shown in the UCSC genome browser. Y-axis shows normalized RPKM counts. **(F)** Expression of genes located within 50kb from any of the breast cancer subtype-selective enhancers defined in Fig. 1E in TNBC and ER+ breast cancers from TCGA (Cancer Genome Atlas Research et al., 2013). Average expression of the gene groups for each patient is summarized in the combined violin and boxplots.

**Supplementary Figure 2.**
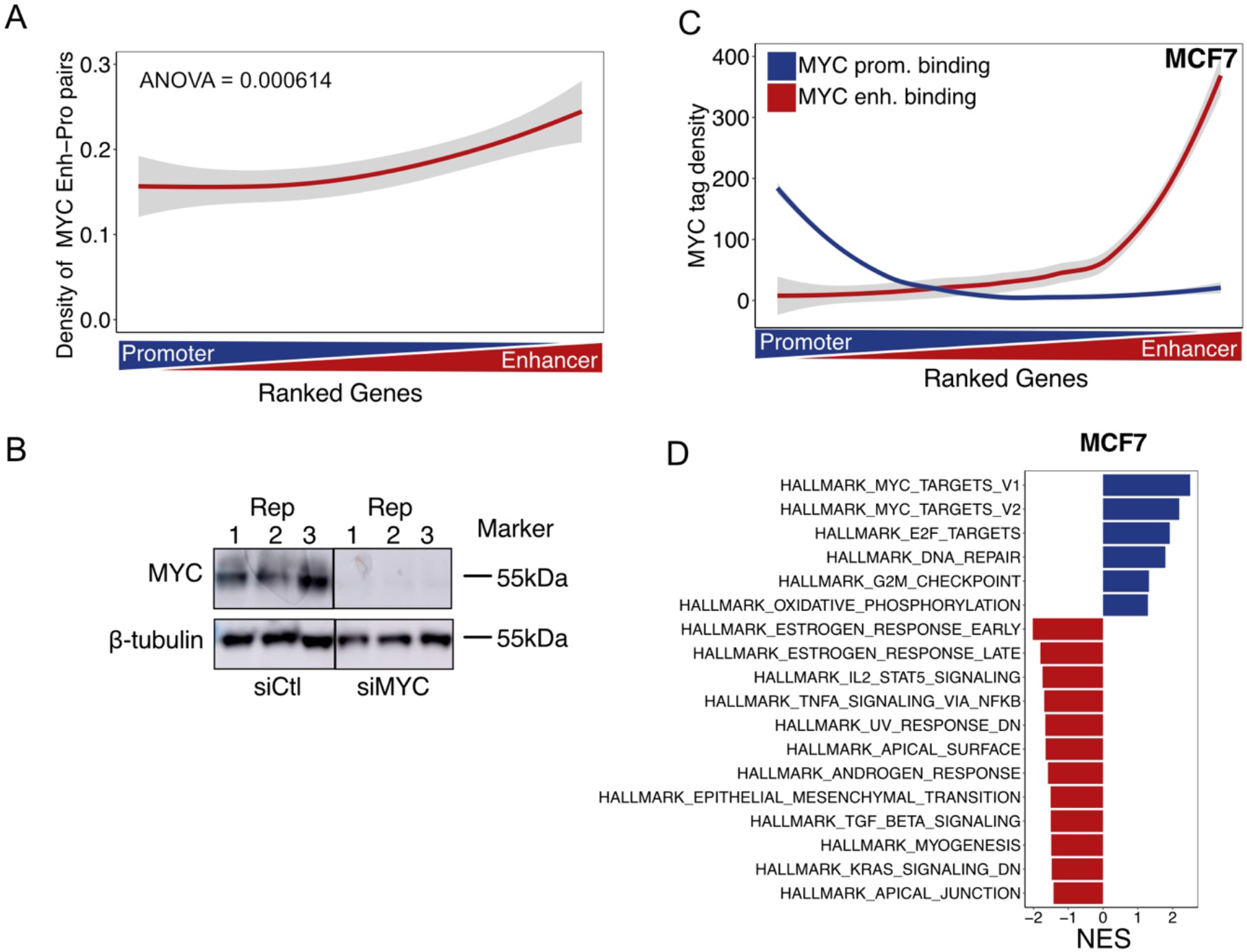
**(A)** Occurrence of enhancer-promoter pairs involving MYC-bound enhancers over the promoter/enhancer ranking of MYC binding in BT549 cells shown in Fig. 2A. Enhancer-promoter pairs were extracted from (Thurman et al., 2012). **(B)** Western blot validation of 72h MYC siRNA knockdown in BT549 cells. (**C-D**) Same as Fig. 2A,C for MYC ChIP-seq in MCF7 cells considering all genes within 50kb of MYC binding sites.

**Supplementary Figure 3.**
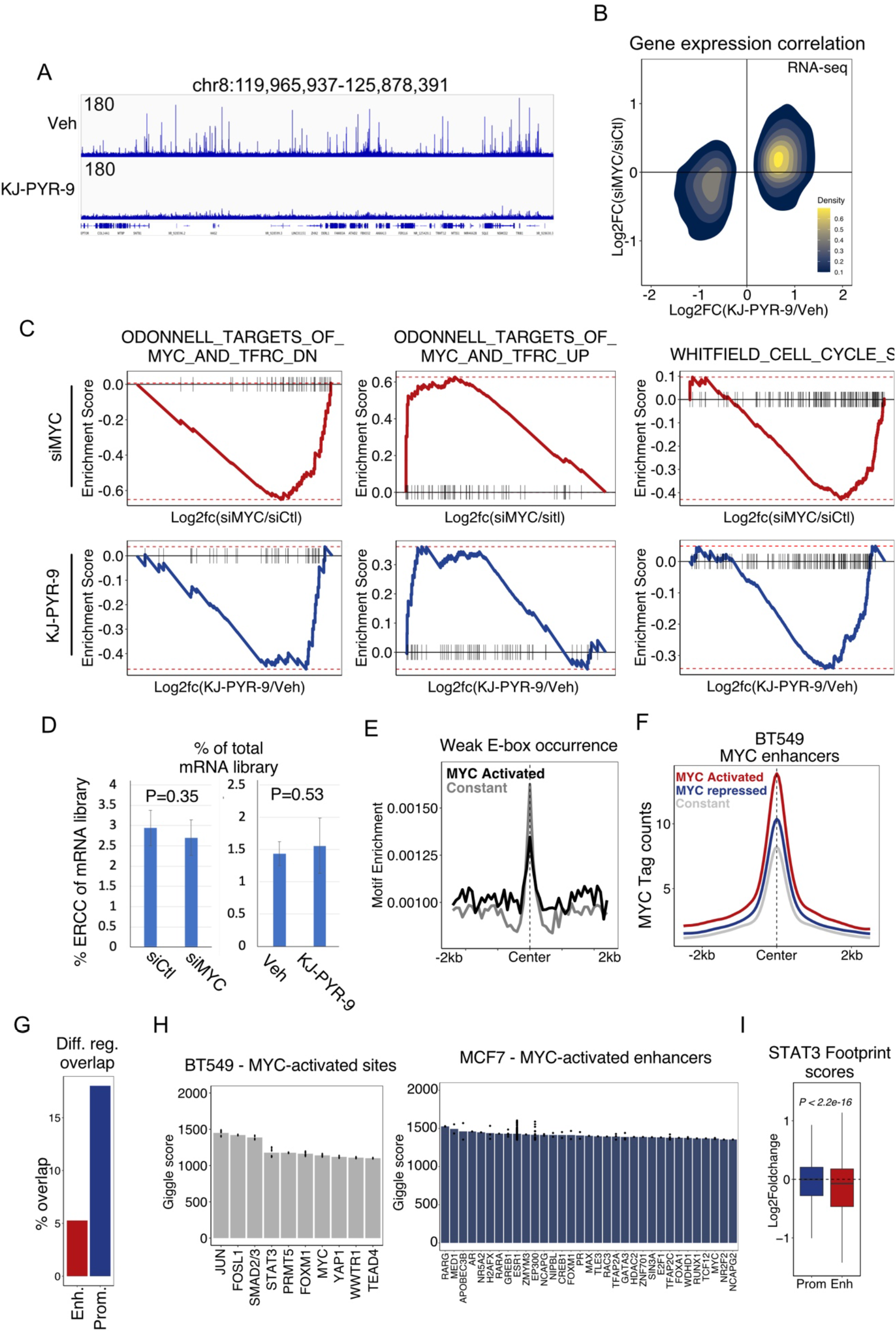
**(A)** UCSC Genome browser screen shot showing MYC ChIP-seq +/- KJ-PYR-9 treatment (3h) for part of chromosome 8. The y-axis shows normalized FPKM counts. **(B)** Differentially expressed genes (Padj<0.05, DEseq2) upon KJ-PYR-9 treatment (3h) in BT549 cells were identified and their response to KJ-PYR-9 was compared with the response to siRNA-mediated knock down of MYC in a scatter plot. The color intensity indicates the density of genes in the scatterplot. **(C)** Enrichment of three MYC target gene sets over genes ranked by log2 fold change in response to KJ-PYR-9 treatment (3h) or siRNA-mediated MYC knock down in BT549 cells. **(D)** Percentage of ERCC spike-ins in RNA-seq libraries from BT549 cells upon siRNA-mediated knock down of MYC or KJ-PYR-9 treatment (3h) (n=3). Error bars represent SEM. P-values were determined by student t-test. **(E)** Enrichment of a weak E-box motif in MYC-bound enhancers significantly activated or constitutively active as determined by MED1 ChIP-seq in response to KJ-PYR-9 treatment (3h) in BT549 cells. **(F)** Average MYC ChIP-seq signal from BT549 cells at MYC-activated, MYC-repressed, or constitutive active enhancers as determined by MED1 ChIP-seq. **(G)** Percentage overlap of the differential MED1 enhancers and promoters upon KJ-PYR-9 treatment (3h) defined in Fig. 3B,C between BT549 and MCF7 cells. **(H)** Giggle analysis (Layer et al., 2018) on MYC-activated enhancers in MCF7 and BT549 cells. A high Giggle score indicates a high overlap with chromatin binding of a given transcription factor as determined by ChIP-seq. Only ChIP-seq performed in triple-negative breast cancer or ER positive breast cancer cell lines were used for this analysis. **(I)** Change in STAT3 footprint scores calculated from corrected cut counts within MYC-bound promoters (n=3811) and enhancers (n=6723) upon KJ-PYR-9 treatment (3h) in BT549 cells. P-value was calculated with Welch’s t-test, two-sided.

**Supplementary Figure 4.**
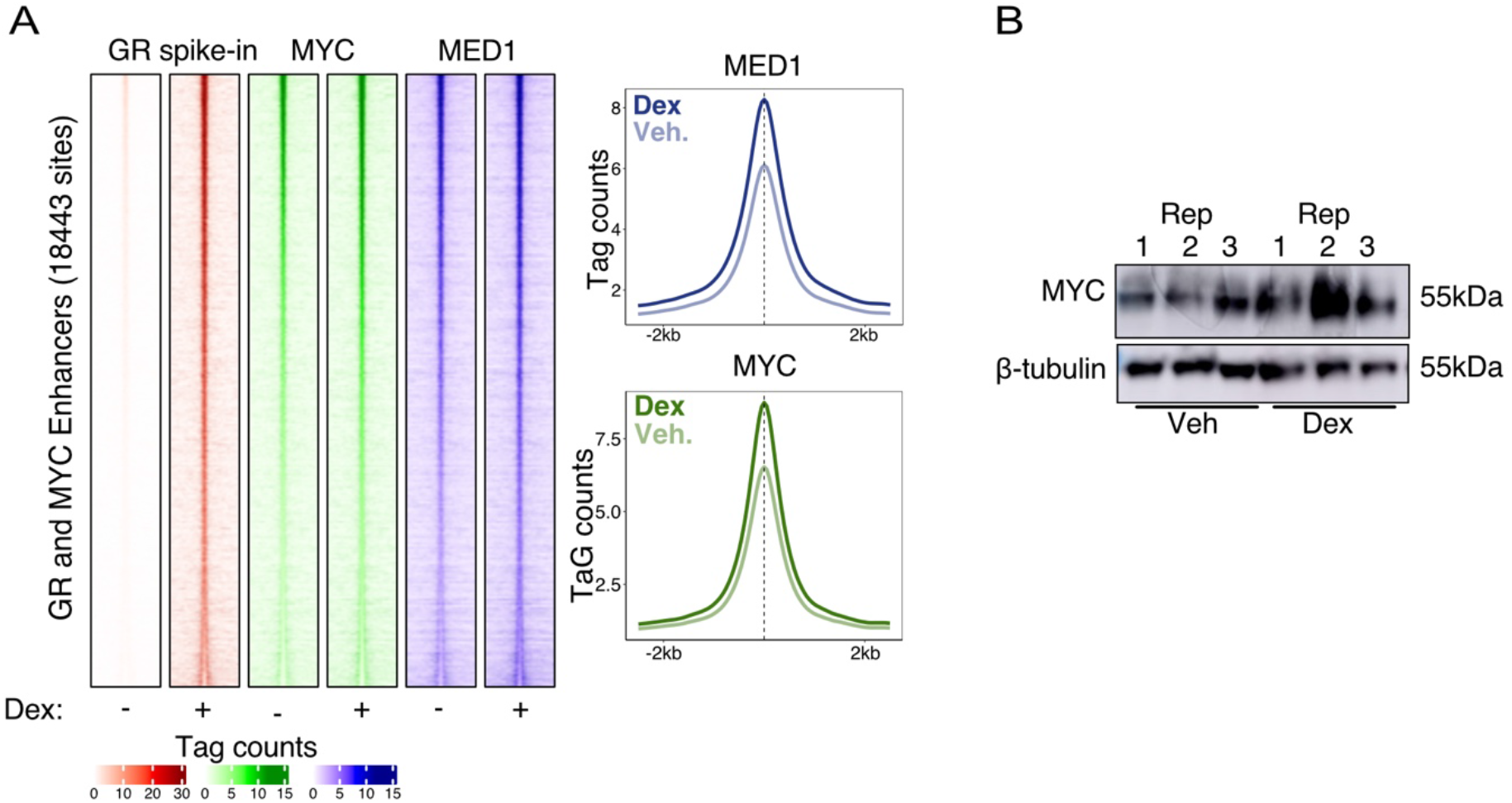
**(A)** Heatmap (left) and average line plots (right) of all common GR and MYC enhancer binding sites upon Dex treatment (2h) in BT549 cells. **(B)** MYC western blot upon Dex treatment (2h) in BT549 cells for three independent biological replicates.

**Supplementary Figure 5.**
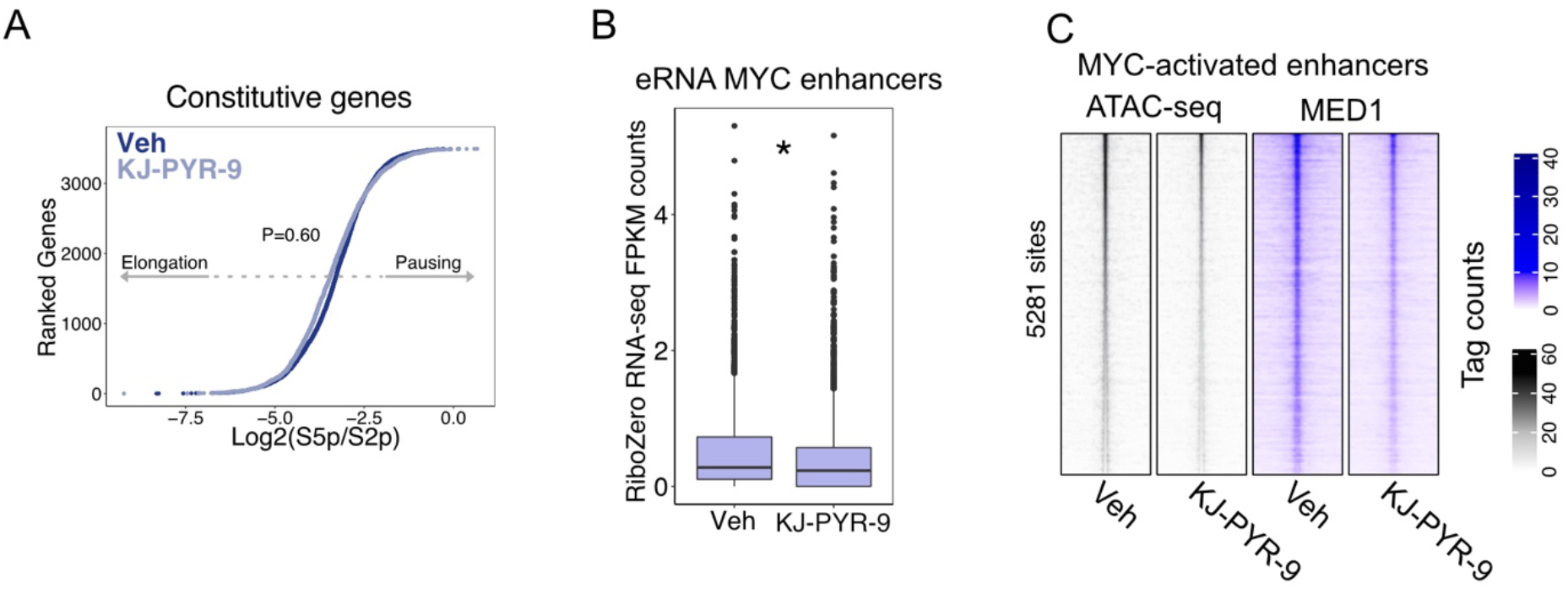
**(A)** Same as Fig. 5A showing genes that are not regulated upon 3h KJ-PYR-9 treatment (Adj. P-value > 0.2) without strong MYC binding at the promoter (Tag count<25) in BT549 cells. P-value was calculated using Welch’s two-sided t-test. **(B)** eRNA within MYC-activated enhancers measured using Ribosomal RNA-depleted (Ribo-Zero) RNA-seq in BT549 cells. * P-value < 0.05 calculated using Welch’s two-sided t-test. **(C)** Chromatin accessibility as determined by ATAC-seq at MYC-activated enhancers in response to KJ-PYR-9 treatment (3h) in BT549 cells.

**Supplementary Figure 6.**
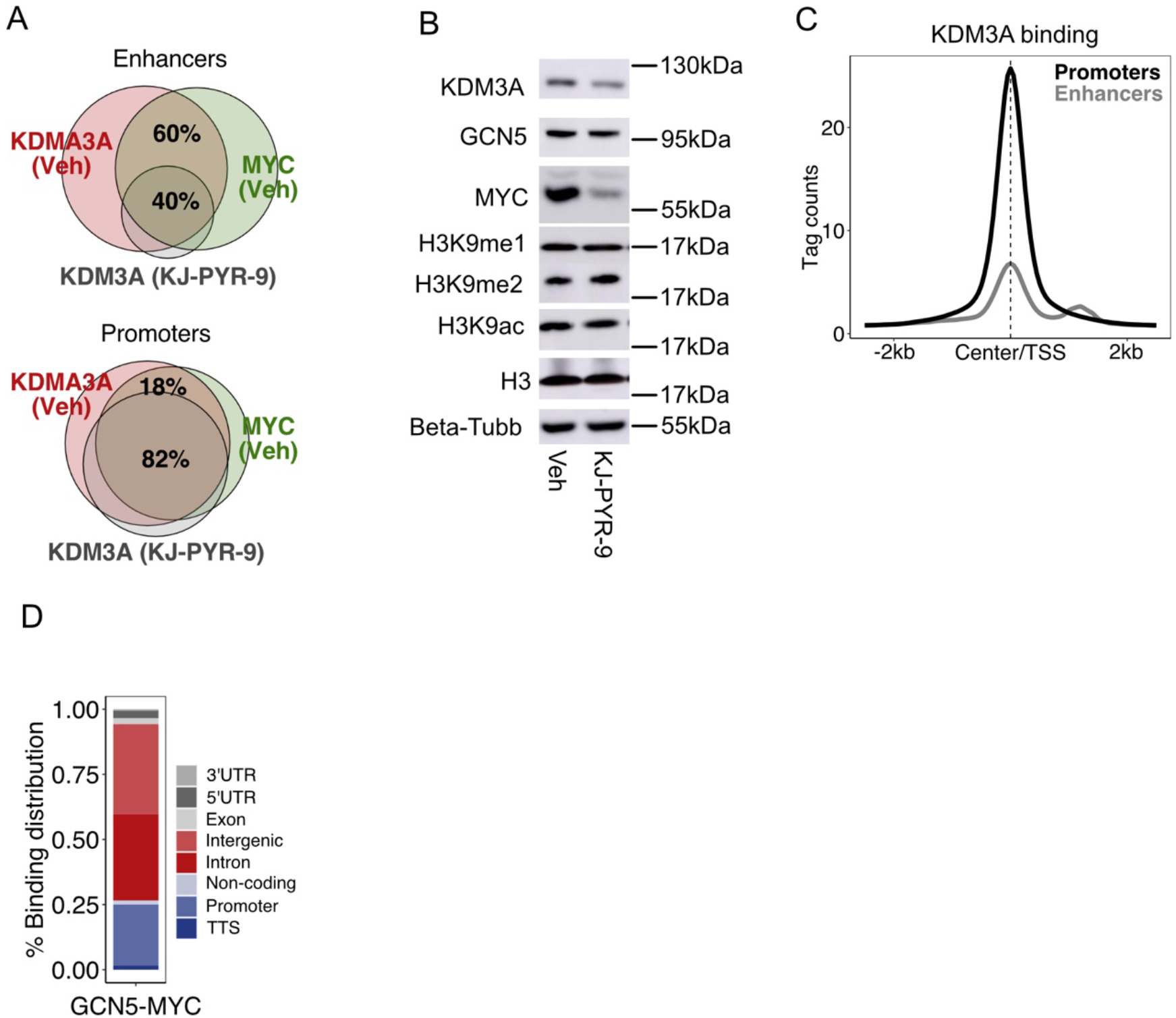
**(A)** Venn diagram showing overlap between KDM3A and MYC binding sites as determined by ChIP-seq in BT549 cells. **(B)** Western blot showing KDM3A, GCN5, MYC, H3K9me1/2, H3K9ac, and H3protein levels upon KJ-PYR-9 treatment (3h) in BT549 cells. β-Tubulin is included as a control. Representative of three independent biological replicates. **(C)** Average plot showing KDM3A binding at promoters versus enhancers as determined by ChIP-seq in BT549 cells. **(D)** Genomic annotation of GCN5 ChIP-seq binding sites overlapping with MYC binding sites.

**Supplementary Figure 7.**
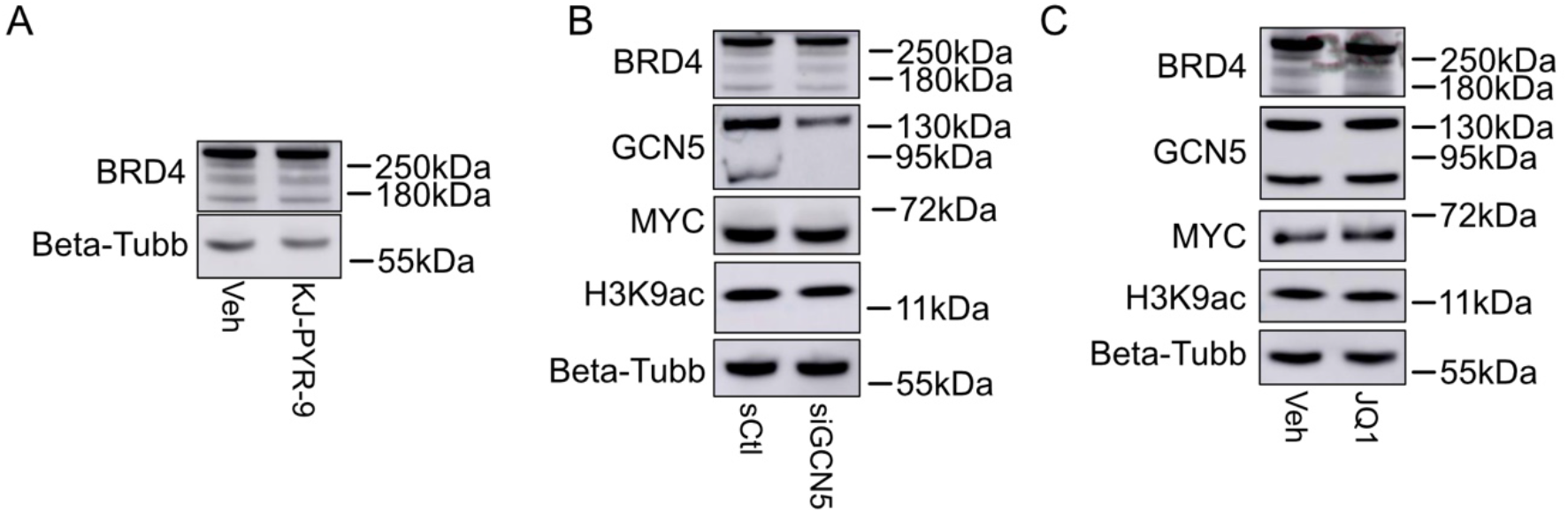
**(A)** Western blot for BRD4 in response to KJ-PYR-9 treatment (3h) in BT549 cells. β-tubulin is included as a control. Representative of two independent biological replicates. **(B)** Western blot showing BRD4, GCN5, MYC, and H3K9ac levels in response to siRNA mediated GCN5 knockdown in BT549 cells. β-tubulin is included as a control. Representative of two independent biological replicates. **(C)** Western blot showing BRD4, GCN5, MYC, and H3K9ac levels upon JQ1 treatment (1h) in BT549 cells. β-tubulin is included as control. Representative of two independent biological replicates.

